# Autocrine IL11 cis-signaling in hepatocytes is an initiating nexus between lipotoxicity and non-alcoholic steatohepatitis

**DOI:** 10.1101/2020.03.11.986802

**Authors:** Jinrui Dong, Eleonora Adami, Sonia P. Chothani, Sivakumar Viswanathan, Benjamin Ng, Wei Wen Lim, Brijesh K. Singh, Jin Zhou, Nicole SJ. Ko, Shamini G. Shekeran, Jessie Tan, Sze Yun Lim, Mao Wang, Pei Min Lio, Paul M. Yen, Sebastian Schafer, Stuart A. Cook, Anissa A. Widjaja

**Author notes:** Correspondence to: Anissa A. Widjaja or Stuart A. Cook, or, 8 College Road 169857, Duke-NUS Medical School, Singapore. These authors jointly supervised this work (Stuart A. Cook, Anissa A. Widjaja). **Author contributions**: S.A.C and A.A.W. conceived and designed the study. J.D, E.A., S.V., B.N., W.W.L., B.K.S., J.T., M.W., and A.A.W. performed *in vitro* cell culture, cell biology and molecular biology experiments. J.D., J.Z., S.G.S, and N.S.J.K. performed *in vivo* studies. S.G.S. and S.Y.L performed histology analysis. J.D., S.P.C., P.M.Y., S.A.C., and A.A.W. analyzed the data. L.P.M and E.A. performed RNA and Ribo sequencing. J.D., E.A., S.S., S.A.C., and A.A.W. prepared the manuscript with input from co-authors.

## Abstract

**Background and aims:** IL11 signaling is important in non-alcoholic steatohepatitis (NASH) but how it contributes to NASH pathologies beyond fibrosis is not known. Here we investigate the role of IL11 signaling in hepatocyte lipotoxicity.

**Methods:** Hepatocytes were stimulated with IL6, IL11, HyperIL6, or HyperIL11 alone or in the presence of soluble gp130 (sgp130) or soluble IL11RA (sIL11RA), or loaded with palmitate in the presence of IgG or anti-IL11RA (X209) antibodies or sgp130. Effects were assessed using colorimetric ALT, GSH, or ELISA assays, immunoblots, and flow cytometry. The relative contributions of IL11 *cis*-versus -*trans* signaling *in vivo* was assessed in two preclinical NASH models using a high fat methionine/choline deficient diet or a Western diet with liquid fructose in C57BL6/Ntac mice injected with AAV8-Alb-Cre, AAV8-Alb-sgp130, in mice with hepatocyte-specific deletion of *Il11ra* (CKO), and in mice with global deletion of *Il11ra* injected with AAV8-Alb-mIl11ra or AAV8-Alb-sIl11ra. Livers and serum were collected; serum samples were analyzed using biochemistry and liver tissues were analyzed by histology, qPCR, immunobloting, hydroxyproline, and GSH assays.

**Results:** We show that lipid-laden hepatocytes secrete IL11, which acts via autocrine *cis-*signaling to cause lipoapoptosis. IL11 causes lipotoxic hepatocyte death through activation of non-canonical signaling pathways and increased NOX4-derived reactive oxygen species. In two preclinical models, hepatocyte-specific deletion of *Il11ra1* protects mice from all aspects of NASH with beneficial effects on body weight. In accordance, restoration of IL11 *cis*-signaling in hepatocytes only in mice globally deleted for *Il11ra1* reconstitutes steatosis and inflammation. Throughout, we found no evidence to support the existence of IL6 or IL11 *trans-*signaling in the liver.

**Conclusion:** We conclude that autocrine IL11-mediated cell death underlies hepatocyte lipotoxicity and that liver fibrosis and inflammation occur subsequently. These data highlight a new disease mechanism for the transition from compensated fatty liver disease to NASH.

## Introduction

Interleukin 11 (IL11) is a fibrogenic factor [1–4] that is elevated in fibrotic precision-cut liver slices across species [5]. IL11 has recently been shown to have negative effects on hepatocyte function after toxic liver insult [6] and, directly or indirectly, contributes to nonalcoholic steatohepatitis (NASH) pathologies [7]. At the other end of the spectrum, a number of earlier publications suggest that IL11 is protective in mouse models of ischemic-, infective- or toxin-induced liver damage [8–13]. However, it is now apparent that the recombinant human IL11 (rhIL11) reagent used in these earlier studies is ineffective in the mouse [6] and the question as to the true biological effect of IL11 in the liver, specifically in hepatocytes, remains open.

IL11 is a member of the interleukin 6 (IL6) cytokine family and, like IL6, binds to its membrane-bound alpha receptor (IL11RA) and glycoprotein 130 (gp130) to signal in *cis*. IL6 itself has been linked to liver function and publications suggest an overall beneficial effect [14–18]. Aside from *cis* signaling, IL6 can bind to soluble IL6 receptor (sIL6R) and signal in *trans*. IL6 *trans* signaling is considered maladaptive in the context of metabolic and autoimmune disease but beneficial for liver regeneration [16]. It is possible that IL11, like IL6, also signals in *trans* but experiments to date have found no evidence for this in tumors or reproductive tissues [19,20].

The factors underlying the transition from non-alcoholic fatty liver disease (NAFLD) to NASH are multifactorial but lipid loading of hepatocytes is of central importance [21]. Certain lipid species are toxic for hepatocytes and lipotoxicity leads to cytokine release causing hepatocyte death along with activation of hepatic stellate cells (HSCs) and immune cells [21,22]. Lipotoxicity, such as that due to palmitate [23], is an early event in NASH and represents a linkage between diet, NAFLD and NASH. While genetic or pharmacological inhibition of IL6 *cis-*signaling worsens steatosis phenotypes [17,18,24], a role for IL11 in hepatic lipotoxicity has not been described.

In the current study, we used a range of *in vitro* and *in vivo* approaches to address key questions regarding IL11 in hepatocyte biology, NAFLD and NASH: (1) Defining the true role of IL11 *cis-* and *trans-*signaling in human hepatocytes, (2) examining whether lipotoxicity is related to IL11 activity in hepatocytes, (3) establishing whether IL11 or IL6 *trans-*signaling contributes to NASH, (4) dissecting the inter-relationship between IL11 *cis-*signaling in hepatocytes and the development of steatosis, hepatocyte death, inflammation, and fibrosis. These studies reveal unexpected aspects of IL6 and IL11 biology and demonstrate an unambiguous pathogenic effect of lipotoxicity-activated, autocrine IL11 *cis-*signaling in hepatocytes that initiates the transition from NAFLD to NASH.

## Methods

### Cell Culture

#### Primary human hepatocytes culture

Primary human hepatocytes (5200, ScienCell) were maintained in hepatocyte medium (5201, ScienCell) supplemented with 2% fetal bovine serum, 1% Penicillin-streptomycin at 37°C and 5% CO2. Hepatocytes (P2-P3) were serum-starved overnight unless otherwise specified in the methods prior to 24 hours stimulation with different doses of various recombinant proteins as outlined in the main text and/or figure legends.

#### Primary mouse hepatocytes culture

Mouse hepatocytes (ABC-TC3928, AcceGen Biotech) were maintained in mouse hepatocyte medium (ABC-TM3928, AcceGen Biotech) supplemented with 1% Penicillin-streptomycin.

#### HepG2 culture

HepG2 (ATCC) were cultured in Eagle’s Minimum Essential Medium (30-2003, ATCC) supplemented with 10% FBS.

#### AML12 culture

AML12 (ATCC) were cultured in DMEM:F12 Medium (30-2006, ATCC) supplemented with 10% FBS, 10 µg/ml insulin, 5.5 µg/ml transferrin, 5 ng/ml selenium, and 40 ng/ml dexamethasone.

#### THP-1 culture

THP-1 (ATCC) were cultured in RPMI 1640 (A1049101, Thermo Fisher) supplemented with 10% FBS and 0.05mM β-mercaptoethanol. THP-1 cells were differentiated with 10 ng/ml of PMA in RPMI 1640 for 48 hours.

### Palmitate (saturated fatty acid) treatment in vitro

Palmitate:BSA conjugated solution in the ratio of 6:1 was prepared as described earlier [25]. Palmitate (0.5 mM) conjugated in fatty acids free BSA was used to treated cells as described in figure legends; 0.5% BSA solution was used as control.

### Flow cytometry

For surface IL11RA, IL6R, and gp130 analysis, primary human hepatocytes and THP-1 cells were stained with IL11RA, IL6R, or gp130 antibody and the corresponding Alexa Fluor 488 secondary antibody. Cell death analysis was performed by staining primary human hepatocytes with Dead Cell Apoptosis Kit with Annexin V FITC and PI (V13242, Thermo Fisher). PI^+ve^ cells were then quantified with the flow cytometer (Fortessa, BD Biosciences) and analyzed with FlowJo version X software (TreeStar): the preliminary FSC/SSC gates of the starting cell population was 10,000 events. Debris (SSC-A vs FSC-A) and doublets (FSC- H vs FSC- A) were excluded. Boundaries between “positive” and “negative” staining were set at 10^3 for PI staining.

### Immunofluorescence (IF)

Primary human hepatocytes were seeded on 8-well chamber slides (1.5×10^4^ cells/well) 24 hours before the staining. Cells were fixed in 4% PFA for 20 minutes, washed with PBS, and non-specific sites were blocked with 5% BSA in PBS for 2 hours. Cells were incubated with IL11RA, IL6R, gp130, or Albumin antibody overnight (4°C), followed by incubation with the appropriate Alexa Fluor 488 secondary antibody for 1 hour. Chamber slides were dried in the dark and 5 drops of mounting medium with DAPI were added to the slides for 15 minutes prior to imaging by fluorescence microscope (Leica).

### Animal models

Animal experiments were performed under the guidelines of SingHealth Institutional Animal Care and Use Committee (IACUC). Mice were maintained in SPF environment and provided with food and water *ad libitum*.

#### Mouse models of metabolic liver disease

1. *HFMCD* 6-8 weeks old C57BL/6N, *Il11ra1*^*-/-*^ mice, and *Il11ra1*^*loxP/loxP*^ and their respective control were fed with methionine- and choline-deficient diet supplemented with 60 kcal% fat (HFMCD, A06071301B16, Research Diets) for 4 weeks. Control mice received normal chow (NC, Specialty Feeds).
2. *WDF* 6-8 weeks old C57BL/6N, *Il11ra1*^*-/-*^ mice, and *Il11ra1*^*loxP/loxP*^ and their respective control were fed western diet (D12079B, Research Diets) supplemented with 15% weight/volume fructose in drinking water (WDF) for 16 weeks. Control mice received NC and tap water.

#### Il11ra1-deleted mice (KO)

6-8-week old male *Il11ra1*^*-/-*^ mice (B6.129S1-*Il11ra*^*tm1Wehi*^/J, Jackson’s Laboratory) were intravenously injected with 4×10^11^ genome copies (gc) of AAV8-*Alb*-*mbIl11ra1* or AAV8-*Alb*-*sIl11ra1* virus to induce hepatocyte specific expression of mouse *Il11ra1* or soluble *Il11ra1*, respectively. As controls, both *Il11ra1*^*-/-*^ mice and their wildtype littermates (*Il11ra1*^*+/+*^) were intravenously injected with 4×10^11^ gc AAV8-Alb-Null virus. 3 weeks after virus injection, mice were fed with HFMCD, WDF, or NC. Durations of diet are outlined in the main text and/or figure legends.

#### In vivo administration of soluble gp130

6-8-week old male C57BL/6N mice (InVivos) were injected with 4×10^11^ gc AAV8-*Alb*-*sgp130* virus to induce hepatocyte specific expression of soluble gp130; control mice were injected with 4×10^11^ gc AAV8-*Alb*-Null virus. 3 weeks following virus administration, mice were fed with HFMCD, WDF, or NC for durations that are outlined in the main text and/or figure legends.

#### Il11ra-floxed mice (CKO)

*Il11ra*-floxed mice, in which exons 4 to 7 of the *Il11ra1* gene were flanked by loxP sites, were created using CRISPR/Cas9 system as previously described[26]. To induce the specific deletion of *Il11ra1* in hepatocytes, 6-8-week old male homozygous *Il11ra1*-floxed mice were intravenously injected with AAV8-*Alb*-Cre virus (4×10^11^gc); a similar amount of AAV8-*Alb*-Null virus were injected into homozygous *Il11ra1*-floxed mice as controls. The AAV8-injected mice were allowed to recover for three weeks prior to HFMCD, WDF, or NC feeding. Knockdown efficiency was determined by Western blotting of hepatic IL11RA.

## Results

### High levels of IL11RA expression in primary human hepatocytes

We first assessed the expression levels of IL6R and IL11RA in healthy human liver by immunohistochemistry. We found robust expression of IL11RA throughout human liver sections but very limited staining of IL6R (**Fig. 1A**). Flow cytometry studies confirmed that IL11RA and gp130 were highly expressed in the majority of primary human hepatocytes (92.6% and 91.9%, respectively) but only few hepatocytes (3.0%) expressed low amounts of IL6R (**Fig. 1B; Supplementary Fig. 1A**). In keeping with this, RNA-seq and Ribo-seq studies showed *IL11RA* and *gp130* transcripts to be highly expressed and actively translated in hepatocytes. In contrast, few *IL6R* transcripts were observed and there was almost no detectable *IL6R* translation (**Fig. 1C-E; Supplementary Fig. 1B and C; Supplemental Info. 1**). Immunofluorescence staining of primary human and mouse hepatocytes, and some of the most commonly used hepatocyte-like cell lines revealed that all of these cells consistently had high IL11RA expression but no detectable IL6R, which was expressed instead on immune cells (**Supplementary Fig. 1D**). We excluded IL6R shedding to the media, where levels were just above the lower limit of detection (**Fig. S1E**). Overall, these data show very low IL6R expression in primary hepatocytes, which implies a limited role for IL6 *cis-*signaling in these cells, but strong co-expression of both IL11RA and gp130.

**Figure 1.**
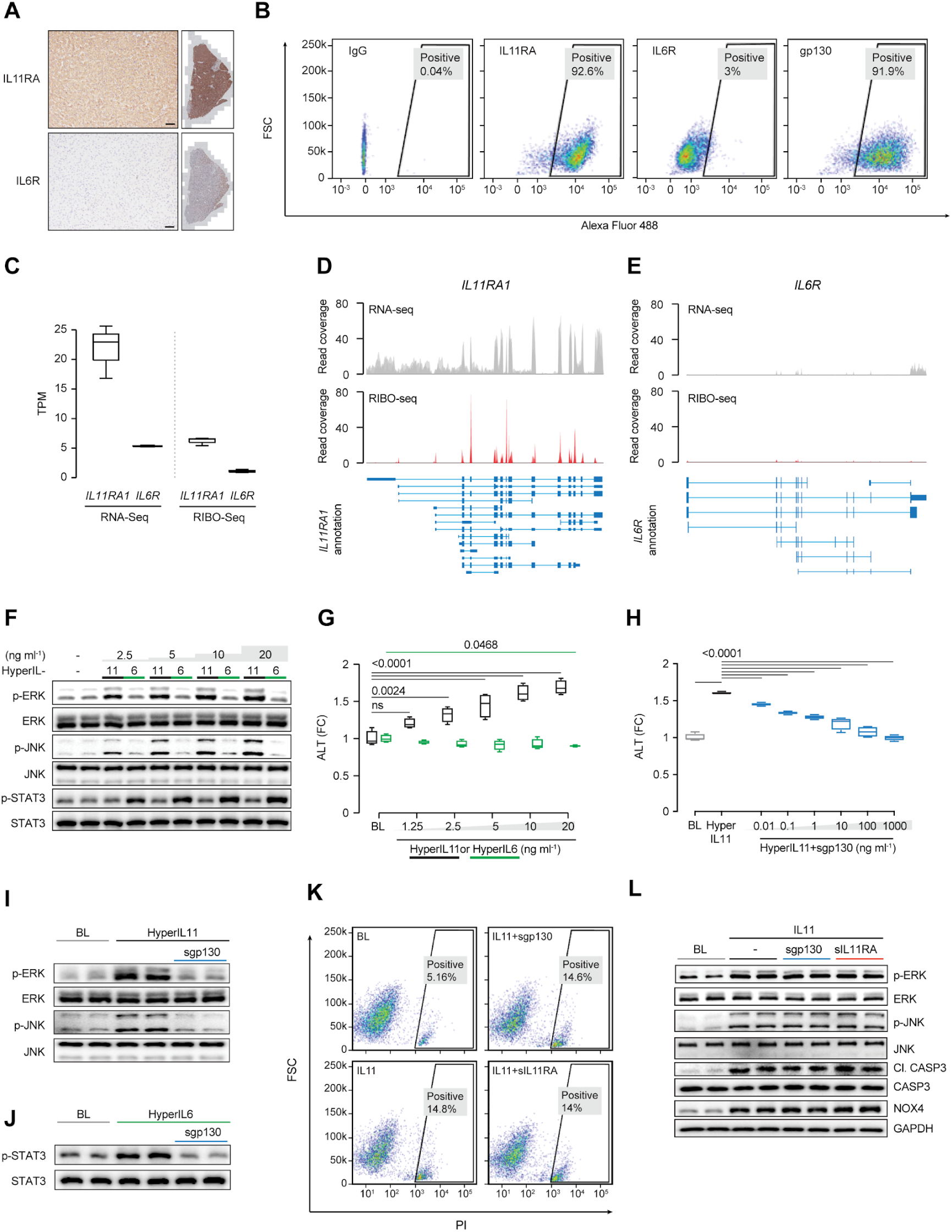
IL11RA is highly expressed in hepatocytes and IL11 cis-signaling is hepatotoxic. (A) Immunohistochemistry staining of IL11RA and IL6R in healthy human liver sections. (B) Representative flow cytometry forward scatter (FSC) and fluorescence intensity plots of IL11RA, IL6R and gp130 staining on hepatocytes. (C) Abundance of *IL11RA1* and *IL6R* reads in hepatocytes at basal based on RNA-seq (left) and Ribo-seq (right) (Transcripts per million, TPM). (D and E) Read coverage of (D) *IL11RA1* and (E) *IL6R* transcripts based on RNA-seq (grey) and Ribo-seq (red) of primary human hepatocytes (n=3). (F and G) (F) Western blots showing ERK, JNK and STAT3 activation status and (G) ALT secretion by hepatocytes following stimulation of either hyperIL11 or hyperIL6 over a dose range. (H) ALT levels in the supernatants of hepatocytes stimulated with hyperIL11 alone or in the presence of increasing amounts of soluble gp130 (sgp130). (I and J) Western blots of hepatocyte lysates showing (I) phosphorylated ERK and JNK and their respective total expression in response to hyperIL11 stimulation alone or with sgp130 and (J) phosphorylated STAT3 and total STAT3 in response to hyperIL6 stimulation with and without sgp130. (K) Representative FSC plots of propidium Iodide (PI) staining of IL11-stimulated hepatocytes in the presence of sgp130 or soluble IL11RA (sIL11RA). (L) Western blots showing p-ERK, p-JNK, cleaved Caspase3 and their respective total expression, NOX4, and GAPDH in hepatocytes in response to IL11 stimulation alone or in the presence of sgp130 or sIL11RA. (B-L) primary human hepatocytes; (F-L) 24 h stimulation; (F-L) hyperIL11, hyperIL6, IL11 (20 ng/ml), sgp130, sIL11RA (1 µg/ml). (C, G-H) Data are shown as box-and-whisker with median (middle line), 25th–75th percentiles (box) and min-max values (whiskers).

### IL11 cis-signaling is a primary driver of hepatocyte death

Given the lack of IL6R in human hepatocytes we needed to employ a synthetic IL6 *trans-*signaling construct (hyperIL6) to activate IL6 signaling and compared this with a synthetic IL11 *trans-*signaling complex (hyperIL11). HyperIL11, like IL11 itself [6], activated ERK and JNK in a dose-dependent manner (2.5 ng/ml to 20 ng/ml). In contrast, IL6 *trans-*signaling did not activate non-canonical signaling pathways but instead dose-dependently induced STAT3 activation (**Fig. 1F**). Thus, IL11 or IL6 with their cognate receptors in pre-formed synthetic complexes activate different intracellular pathways when bound to gp130 on hepatocytes, which is a novel and intriguing finding. HyperIL11 caused a dose-dependent increase in alanine transaminase (ALT) in the media of primary human hepatocyte cell cultures whereas hyperIL6 (20 ng/ml) had a significant, albeit limited, protective effect (ALT fold change (FC)=0.9; P=0.0468) (**Fig. 1G**). Soluble gp130 (sgp130) is a selective inhibitor of trans-signaling complexes acting through gp130 [16]. Consistent with its reported decoy effects, sgp130 blocked the activation of signaling pathways downstream of both hyperIL11 (p-ERK/p-JNK) and hyperIL6 (p-STAT3) and also inhibited the hepatotoxic effects of hyperIL11 (**Fig. 1H-J**).

We then probed for the existence of physiological IL11 *trans-*signaling. We first stimulated cells with IL11 in the presence of either soluble gp130 (sgp130, to inhibit putative *trans-*signaling) or soluble IL11RA (sIL11RA, to potentiate putative *trans-* signaling). IL11-induced Caspase3-dependent hepatocyte death, NOX4 upregulation, ERK and JNK signaling were unaffected by sgp130 or sIL11RA (**Fig. 1 and L; Supplementary Fig. 2A**). Furthermore, IL11 dose-dependently (0.625 ng/ml to 20 ng/ml) caused hepatocyte cell death, which was unaffected by the addition of sgp130 (1 µg/ml) or sIL11RA (1 µg/ml) (**Supplementary Fig. 2B**). Reciprocally, increasing doses of sgp130 or sIL11RA had no effect on ALT release from IL11-stimulated hepatocytes (**Supplementary Fig. 2C**). These data argue against the existence of IL11 *trans-* signaling in a biological context.

### IL11 cis-signaling underlies lipotoxicity in hepatocytes

To begin to examine the role of IL11 in fatty liver disease, we modelled hepatocyte lipotoxicity, viewed as an initiating pathology for NASH and related to cytokine release [21]. To do so, we loaded hepatocytes with palmitate using a concentration of saturated fatty acids (sFAs) seen in the serum of NAFLD patients [27]. Compared to control, sFA loaded cells secreted greater amounts of IL11 (FC=28, P<0.0001) and also IL6, CCL2 and CCL5 (**Fig. 2A-D**). Lipid loaded hepatocytes underwent apoptotic cell death and also necrotic release of ALT (**Fig. 2E-G**).

**Figure 2.**
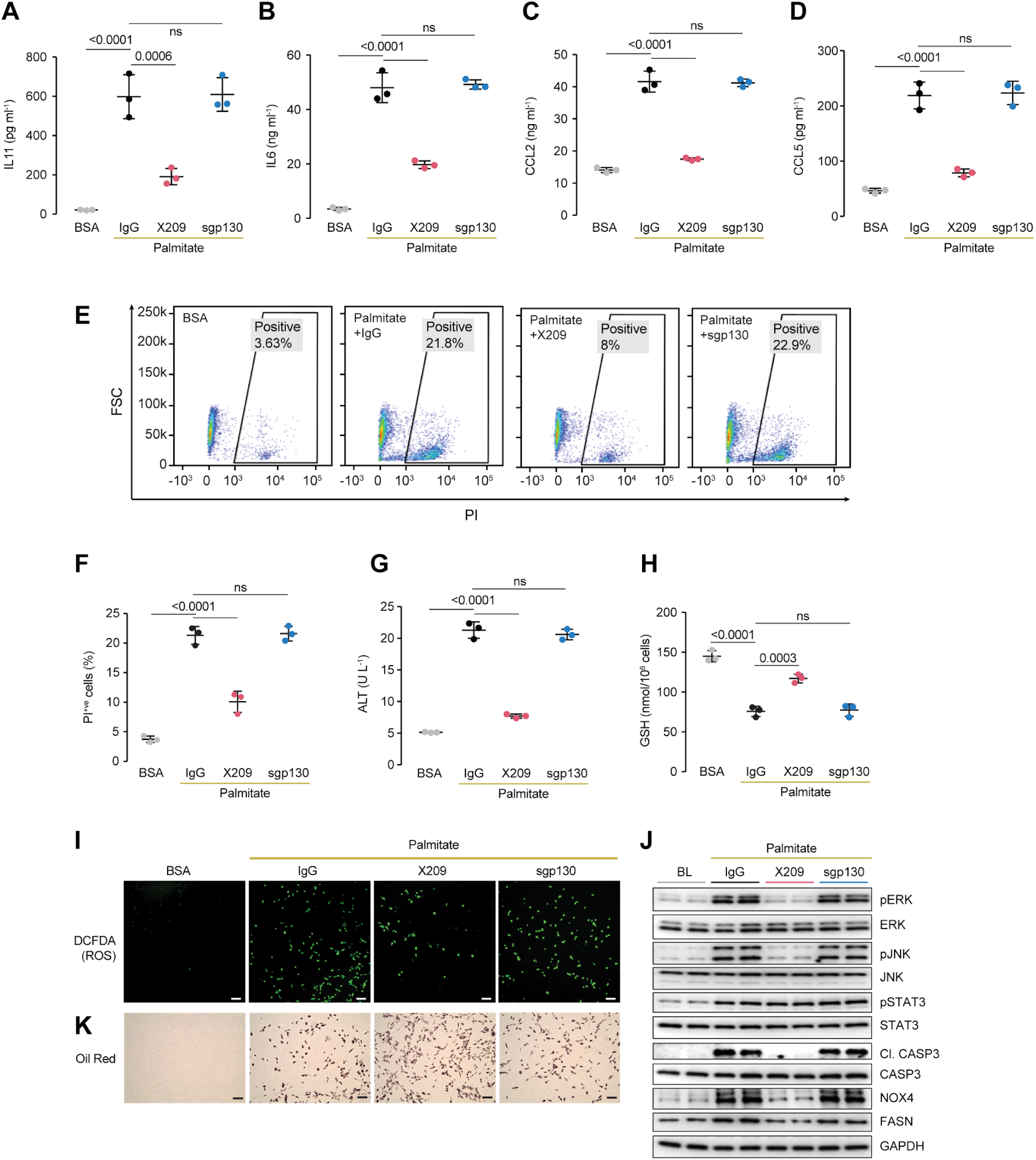
Lipid laden hepatocytes secrete IL11 that drives lipotoxic phenotypes through autocrine activation of non-canonical IL11 *cis*-signaling. (A-K) Data for palmitate loading experiment on primary human hepatocytes in the presence of either IgG (2 µg/ml), anti-IL11RA (X209, 2 µg/ml), or sgp130 (1 µg/ml). (A) IL11, (B) IL6, (C) CCL2, and (D) CCL5 protein secretion levels as measured by ELISA of supernatants. (E and F) (E) Representative FSC plots and (F) quantification of PI^+ve^ hepatocytes stimulated with palmitate. (G) ALT levels in supernatants. (H) Hepatocyte glutathione (GSH) levels. (I) Representative fluorescence images of DCFDA (2’, 7’-dichlorofluorescein diacetate) staining for ROS detection (scale bars, 100 µm). (J) Western blots of pERK, ERK, pJNK, JNK, cleaved Caspase3, Caspase3, NOX4, FASN and GAPDH. (K) Representative images of Oil Red O staining (scale bars, 100 µm). (A-D, F-H) Mean±SD; Tukey-corrected Student’s *t*-test.

To test if IL11 secretion from lipid laden hepatocytes was mechanistically related to lipotoxicity in a *cis-* or *trans-*signaling manner, we incubated cells with either anti-IL11RA antibody (X209) or sgp130. X209 reduced the secretion of all cytokines, including IL11 itself, whereas sgp130 had no effect (**Fig. 2A-G**). This suggests an autocrine loop of IL11 *cis-*signaling underlies hepatocyte lipotoxicity. Using hyperIL11 stimulation, which is not detected by IL11 enzyme-linked immunosorbent assay [1], we experimentally established the existence of autocrine IL11 signalling in hepatocytes (**Supplementary Fig. 2D**). The production of reactive oxygen species (ROS) from damaged mitochondria is important for lipotoxicity [22] and ROS from NOX4 is also pertinent in NASH [28]. We found that X209 prevented ROS production in sFA-loaded hepatocytes and that this was accompanied by partial restoration of glutathione (GSH) levels (**Fig. 2H and I**).

We then studied signaling events. Lipotoxicity is strongly associated with activation of JNK, which drives caspase-3 activation and lipoapoptosis. Accordingly, palmitate loaded hepatocytes exhibited JNK activation and caspase-3 cleavage, as well as ERK phosphorylation (**Fig. 2J**). This pattern was notably similar to the effects seen with IL11 stimulation (**Fig. 1F**). X209 inhibited palmitate-induced signaling events as well as fatty acid synthase (FASN) upregulation and caspase3 activation despite similar sFA uptake by hepatocytes (**Fig. 2J and K**). NOX4 was upregulated by palmitate and also inhibited by X209 (**Fig 2J**). While STAT3 was activated by sFA loading, this effect is independent of IL11RA-mediated signaling and unrelated to lipoapoptosis (**Fig. 2J**). Throughout these experiments sgp130 had no effect. These data show that palmitate-induced IL11 secretion and autocrine, feed-forward IL11 *cis*-signaling is an important mechanism for hepatocyte lipotoxicity.

### No evidence for IL11 or IL6 *trans*-signaling in two NASH models

We then tested whether *trans-*signaling underlies NASH *in vivo* using two preclinical mouse NASH models: The Western Diet supplemented with fructose (WDF) model and the methionine- and choline-deficient high fat diet (HFMCD) model. The WDF model is associated with obesity, hyperlipidemia and insulin resistance and seen as translatable to common forms of human NASH, as in diabetic patients. The HFMCD model stimulates rapid onset NASH, specifically driven by hepatocyte lipotoxicity, that is associated with weight loss in the absence of insulin resistance. Lipotoxicity is common to both models whereas obesity and insulin resistance are not.

Three weeks prior to starting either the WDF or HFMCD diet, mice were injected with an AAV8 virus encoding either albumin promoter-driven sgp130 (AAV8-Alb-sgp130), which contains the whole extracellular domain of mouse gp130 protein (amino acid 1 to 617), or albumin promoter alone (AAV8-Alb-Null) (**Fig. 3A, Supplementary Fig. 3A, and 4A**). AAV8-Alb-sgp130 administration induced high levels of sgp130 in the liver, which was also detectable in the peripheral circulation, suitable for both local and systemic inhibition of putative IL6 or IL11 *trans-*signaling (**Fig. 3B, Supplementary Fig. 3B, 4B, and C**).

**Figure 3.**
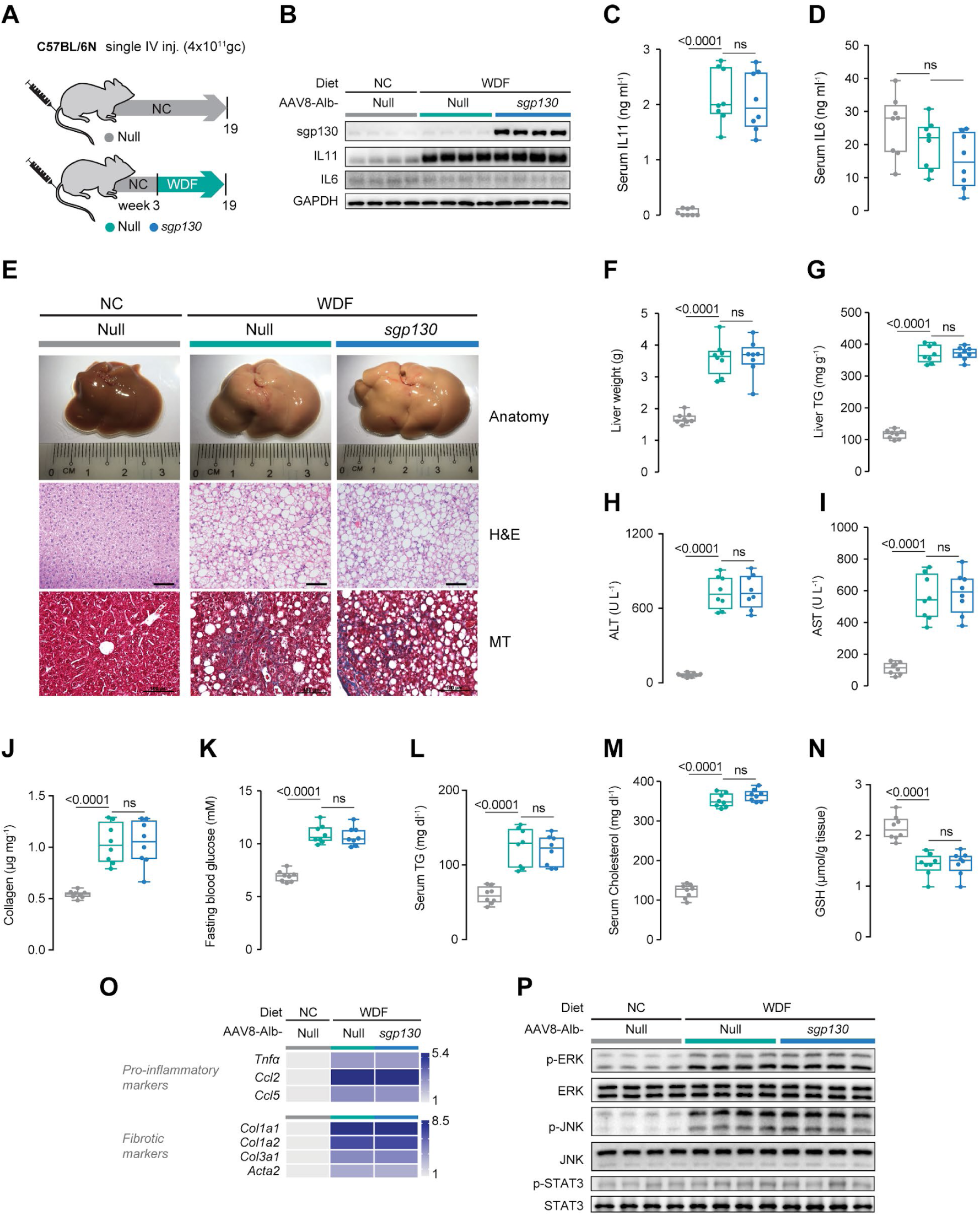
Inhibition of IL6 family cytokine *trans*-signaling has no effect on NASH or metabolic phenotypes in mice on Western Diet supplemented with fructose. (A) Schematic of WDF feeding in mice with hepatocyte-specific expression of sgp130 for data shown in (B-P). Three weeks following AAV8-Alb-Null or AAV8-Alb-sgp130 virus injection, mice were fed WDF for 16 weeks. (B) Western blots showing hepatic levels of sgp130, IL11, IL6, and GAPDH as internal control. (C) Serum IL11 levels. (D) Serum IL6 levels. (E) Representative gross anatomy, H&E-stained (scale bars, 50 µm) and Masson’s Trichrome (scale bars, 100 µm) images of livers. (F) Liver weight. (G) Hepatic triglycerides content. (H) Serum ALT levels. (I) Serum AST levels. (J) Hepatic collagen levels. (K) Fasting blood glucose levels. (L) Serum triglycerides levels. (M) Serum cholesterol levels. (N) Hepatic GSH content. (O) Hepatic pro-inflammatory and fibrotic genes expression heat map (values are shown in **Supplementary Fig. 3 and E**). (P) Western blots of hepatic p-ERK, ERK, p-JNK, JNK, p-STAT3, and STAT3. (C-N) Data are shown as box-and-whisker with median (middle line), 25th–75th percentiles (box) and min-max values (whiskers), Tukey-corrected Student’s *t*-test.

After 16 weeks of WDF, IL11 levels were strongly upregulated in the liver and the periphery but IL6 expression was unaffected (**Fig. 3B and C**). Mice on WDF became obese (**Supplementary Fig. 3C**), had an approximate 2-fold increase in liver mass and developed severe steatosis and fibrosis by gross morphology, quantitative analysis of liver triglycerides, and histology (**Fig. 3E-G**). These phenotypes were unaffected by high levels of sgp130 expression (**Fig. 3B-G**). Similarly, mice on WDF had elevated levels of ALT, AST, collagen and peripheral cardiovascular risk factors (fasting blood glucose, serum triglycerides and serum cholesterol), along with depleted levels of GSH but none of these parameters were affected by sgp130 (**Fig. 3H-N**). Livers from mice on WDF diet for 16 weeks showed increased expression of pro-inflammatory and fibrosis genes and this signature was unaffected by sgp130-mediated inhibition of putative *trans*-signaling (**Fig. 3O; Supplementary Fig. 3D and E**).

In a second set of experiments we induced NASH using the HFMCD diet (**Supplementary Fig. 4A**). HFMCD diet increased IL11 levels in liver and serum, whereas IL6 levels were slightly lower in the liver and were mildly increased in the periphery (**Supplementary Fig. 4B, 4D, and E**). Mice on HFMCD diet developed rapid and profound steatosis by gross morphology, histology, and molecular assays, which was unaltered by sgp130 expression (**Supplementary Fig. 4F and G**). Hepatocyte damage markers (ALT and AST) were elevated and GSH depleted by HFMCD diet, irrespective of sgp130 expression (**Supplementary Fig. 4H-J**). Similarly, HFMCD-induced liver fibrosis was unchanged by sgp130 expression (**Supplementary Fig. 4F and K**). At the RNA level, the HFMCD diet was associated with dysregulated expression of inflammation and fibrosis genes and these molecular phenotypes were unaffected by sgp130 expression (**Supplementary Fig. 4L and M)**.

At the signaling level, both WDF and HFMCD diets stimulated ERK and JNK activation, consistent with elevated IL11 *cis*-signaling (**Fig. 3P; Supplementary Fig. 4N**). In contrast, pSTAT3 levels in the liver were not elevated by WDF (**Fig. 3P**) and appeared mildly elevated in mice on the HFMCD diet (**Supplementary Fig. 4N**). In all instances, there was no effect of sgp130 on diet-induced signaling events. Overall, these data suggest that neither IL6 nor IL11 *trans*-signaling plays a role in NASH, which is consistent with other studies where IL6 family *trans*-signaling has not been detected [19,20,29,30].

### Hepatocyte-specific IL11 *cis*-signaling is required to initiate NASH

While we found no evidence to support IL11 *trans*-signaling in NASH models, our data suggested increased pathological IL11 signaling in hepatocytes, presumed in *cis*. To test this premise, we administered AAV8-Alb-Cre to *Il11ra1*^*loxP/loxP*^ mice to delete *Il11ra1* in hepatocytes only (CKO mice). CKO mice were then fed either normal chow (NC), HFMCD diet or WDF (**Fig. 4A and 5A**). Liver IL11RA protein was greatly diminished in the CKOs following AAV8-Alb-Cre, showing the model to be effective and suggesting that hepatocytes are the largest hepatic reservoir of Il11ra1 (**Fig. 4B and 5B**).

**Figure 4.**
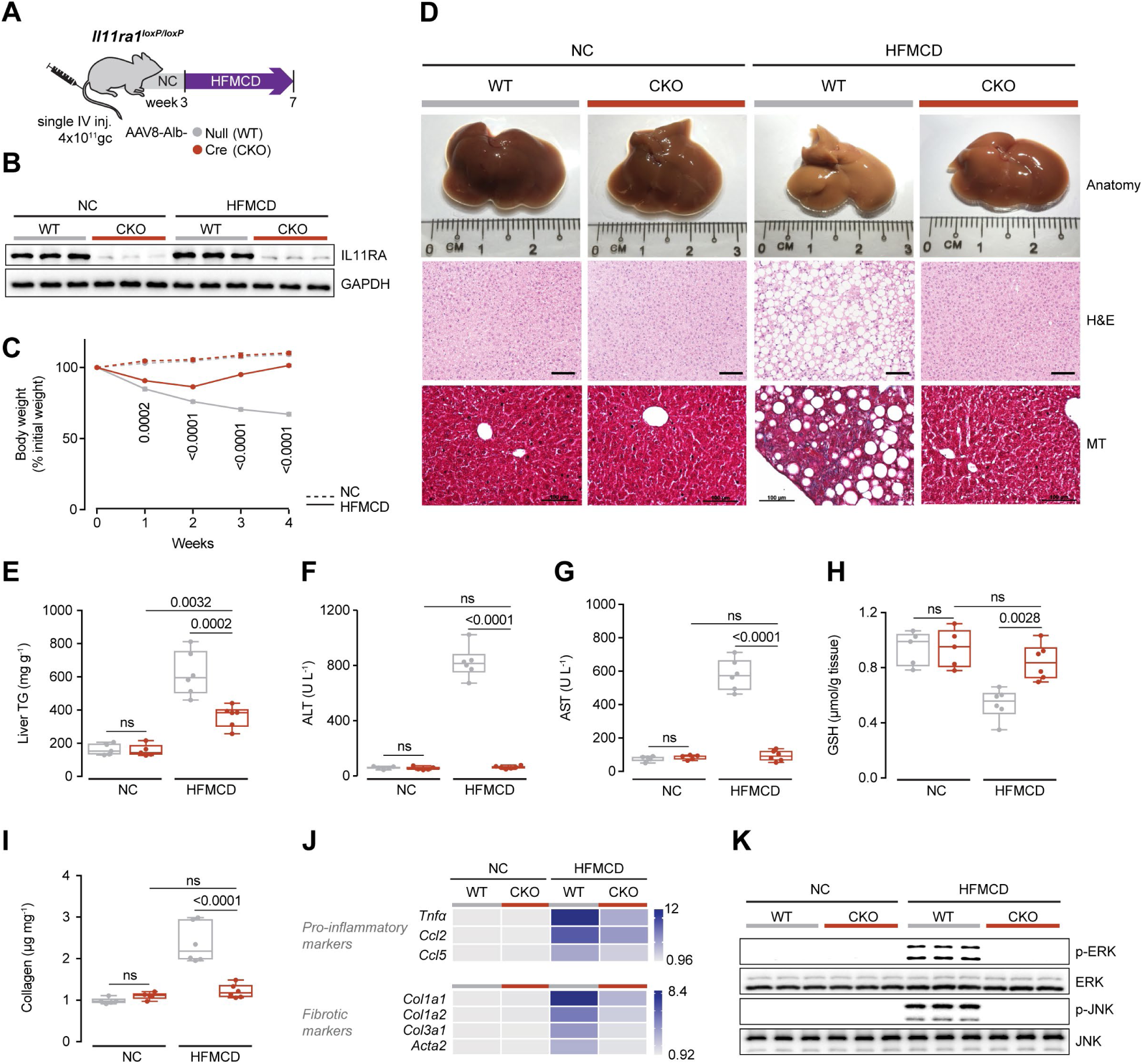
Hepatocyte-specific inhibition of IL11 *cis*-signaling are protected against HFMCD diet-induced cachexia and NASH. (A) Schematic of HFMCD feeding regimen for AAV8-Alb-Cre injected *Il11ra1*^*loxP/loxP*^ (conditional knockout; CKO) mice for experiments shown in (B-K). *Il11ra1*^*loxP/loxP*^ mice were intravenously injected with either AAV8-Alb-Null or AAV8-Alb-Cre to delete *Il11ra1* specifically in hepatocytes three weeks prior to the start of HFMCD diet. (B) Western blots of hepatic IL11RA and GAPDH. (C) Body weight (shown as a percentage (%) of initial body weight). (D) Representative gross anatomy, H&E-stained (scale bars, 50 µm) and Masson’s Trichrome (scale bars, 100 µm) images of livers. (E) Hepatic triglycerides content. (F) Serum ALT levels. (G) Serum AST levels. (H) Hepatic GSH content. (I) Hepatic collagen levels. (J) Heatmap showing hepatic mRNA expression of pro-inflammatory markers (*Tnfα, Ccl2, Ccl5*) and fibrotic markers (*Col1a1, Col1a2, Col3a1, Acta2)*. Values are shown in **Supplementary Fig. 5A and B**. (K) Western blots showing hepatic ERK and JNK activation status. (C) Data are shown as mean ± SEM, 2-way ANOVA with Tukey’s multiple comparison test, statistical significance is shown as the P values between HFMCD WT and CKO; (E-I) Data are shown as box-and-whisker with median (middle line), 25th–75th percentiles (box) and min-max values (whiskers), Sidak-corrected Student’s *t*-test.

**Figure 5.**
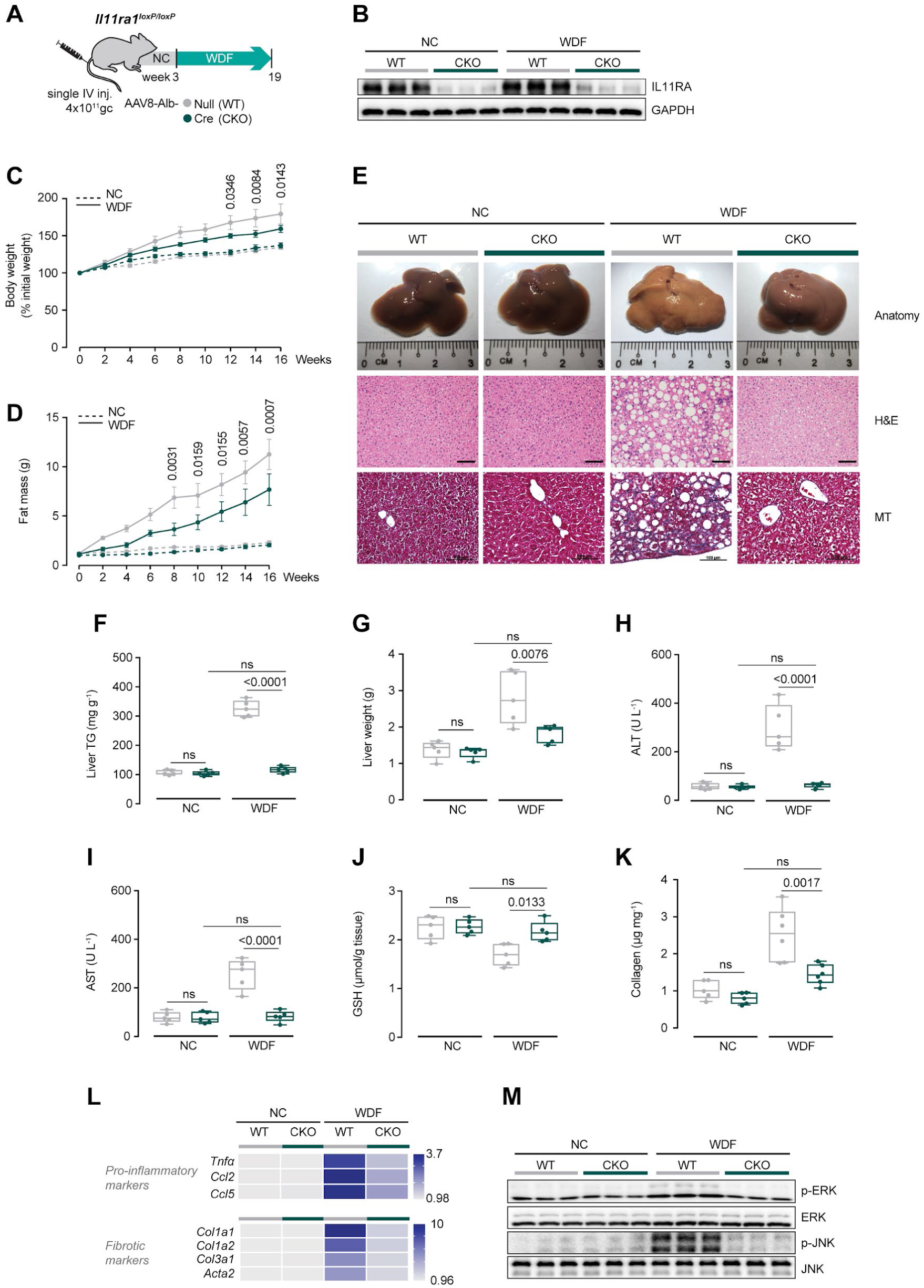
Mice with hepatocyte-specific inhibition of IL11 *cis*-signaling are protected against WDF-induced obesity and NASH. (A) Schematic of WDF-fed control and CKO mice for data shown in (B-M). Three weeks following AAV8-Alb-Null or AAV8-Alb-Cre virus injection, CKO mice were fed WDF for 16 weeks. (B) Western blots showing hepatic levels of IL11RA and GAPDH. (C) Body weight (shown as a percentage (%) of initial body weight). (D) Fat mass. (E) Representative gross anatomy, H&E-stained (scale bars, 50 µm) and Masson’s Trichrome (scale bars, 100 µm) images of livers. (F) Hepatic triglycerides content. (G) Liver weight. (H) Serum ALT levels. (I) Serum AST levels. (J) Hepatic GSH content. (K) Hepatic collagen levels. (L) Hepatic pro-inflammatory and fibrotic genes expression on heat map (values are shown in **Supplementary Fig. 6D and E**). (M) Western blots showing activation status of hepatic ERK and JNK. (C and D) Data are shown as mean ± SEM, 2-way ANOVA with Tukey’s multiple comparison test, statistical significance is shown as the P values between WDF WT and CKO; (F-K) Data are shown as box-and-whisker with median (middle line), 25th–75th percentiles (box) and min-max values (whiskers), Sidak-corrected Student’s *t*-test.

In addition to rapidly stimulating lipotoxicity-driven NASH [31] the HFMCD diet causes weight loss [31]. Surprisingly, weight loss in mice on the HFMCD diet was initially limited and later reversed in CKO mice (**Fig. 4C)**. Mice on WDF gained weight and fat mass throughout the experimental period, as expected. However, and equally surprising, these obesity phenotypes were mitigated in CKO mice (**Fig. 5C and 5D**). These data suggest that inhibition of IL11 signaling is permissive for weight homeostasis, with context-specific anti-cachectic or anti-obesity effects, which requires further study.

By gross morphology, histology and quantitative triglyceride analysis, the CKO mice on either HFMCD or WDF diet were robustly protected from steatosis (**Fig. 4D, 4E, 5E, 5F**) and those on WDF had less hepatomegaly (**Fig. 5G**). Liver damage markers were markedly reduced in CKO mice fed with either HFMCD diet (reduction: ALT, 99%; AST, 97%; P<0.0001 for both) or WDF (reduction: ALT, 98%; AST, 98%; P<0.0001 for both) and found to be comparable to NC control levels (**Fig. 4F, 4G, 5H, 5I**). In both models, GSH levels were diminished in control mice on the NASH diets but normalized in CKOs (**Fig. 4H and 5J**).

Liver fibrosis was greatly reduced in CKO mice on either NASH diet as compared to controls (reduction: HFMCD, 87%; WDF, 64%; P<0.001 for both) (**Fig. 4D, 4I, 5E, 5K**). Upregulation of pro-inflammatory and fibrosis genes in mice on either the HFMCD or WDF diets was also diminished in the CKOs (**Fig. 4J and 5L; Supplementary Fig. 5A, 5B, 6A, 6B**). This suggests that transformation of HSCs to myofibroblasts and activation of immune cells are, in part, secondary to upstream, IL11-driven events in hepatocytes.

Mice on WDF also develop hyperglycemia, hypertriglyceridemia, and hypercholesterolemia, all of which were improved in the CKOs, suggesting an important role for hepatocyte-specific IL11 signalling for NASH phenotypes more generally (**Supplementary Fig. 6C-E**). At the signaling level, both HFMCD diet and WDF resulted in elevated ERK and JNK phosphorylation. This was prevented in CKO mice, consistent with inhibition of IL11 signaling in hepatocytes (**Fig. 4K and 5M**).

### Reconstitution of IL11 *cis-*signaling in hepatocytes alone in *IL11ra1* null mice restores steatohepatitis but not liver fibrosis

To complement our loss-of-function experiments using the CKO mice we employed *in vivo* gain-of-function experiments. To do so, we assessed whether restoring IL11 *cis-* or *trans-*signaling specifically in hepatocytes in mice with global *Il11ra1* deletion (*Il11ra1*^*-/-*^ knockouts (KOs)) resulted in disease. KO mice were injected with AAV8 encoding either the full length, membrane bound *Il11ra1* (*mbIl11ra1*; to reconstitute *cis-*signaling) or a secreted/soluble form of *Il11ra1* (*sIl11ra1*, which constitutes the extracellular portion of *Il11ra1*; to enable *tra*ns-signaling) or a control construct and the animals were then fed with NC, HFMCD diet or WDF (**Fig. 6A; Supplementary Fig. 7A and 8A**).

**Figure 6.**
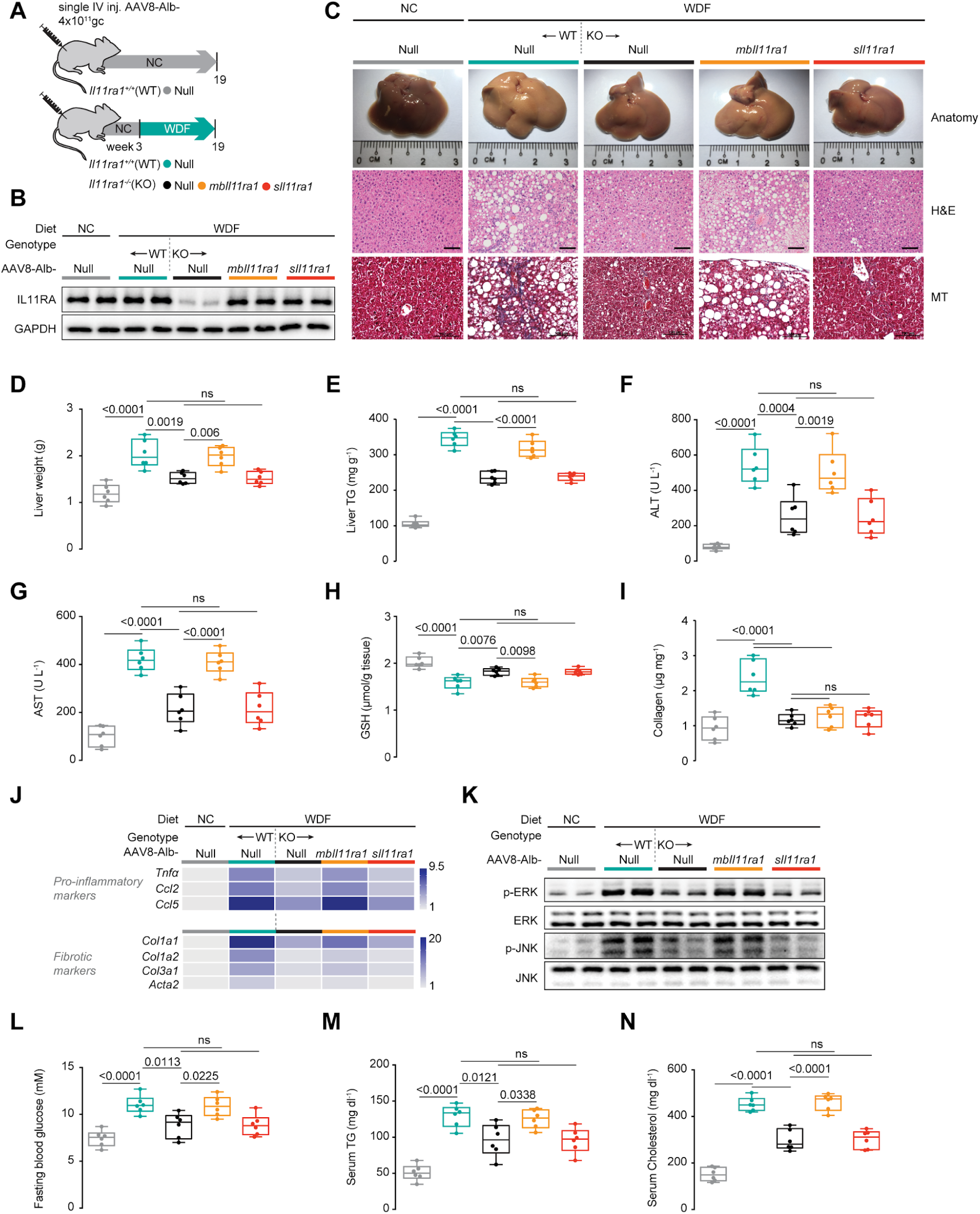
Hepatocyte-specific IL11 *cis*-signaling but not IL11 *trans*-signaling drives steatohepatitis in mice on WDF. (A) Schematic showing WDF feeding regimen of *Il11ra1*^*+/+*^ (WT) and *Il11ra1*^*-/-*^ (KO) mice for experiments shown in (B-N). AAV8-Alb-Null, AAV8-Alb-mbIl11ra1 (full length membrane-bound Il11ra1), and AAV8-Alb-sIl11ra1 (soluble form of Il11ra1) -injected KO mice were given 16 weeks of WDF feeding, three weeks following virus administration. (B) Western blots showing hepatic levels of IL11RA and GAPDH. (C) Representative gross anatomy, H&E-stained (scale bars, 50 µm) and Masson’s Trichrome (scale bars, 100 µm) images of livers. (D) Liver weight. (E) Hepatic triglycerides content. (F) Serum ALT levels. (G) Serum AST levels. (H) Hepatic GSH content. (I) Hepatic collagen content. (J) Hepatic pro-inflammatory and fibrotic genes expression heat map (values are shown in **Supplementary Fig. 7C and D**). (K) Western blots showing activation status of hepatic ERK and JNK. (L) Fasting blood glucose levels. (M) Serum triglycerides levels. (N) Serum cholesterol levels. (D-I, L-N) Data are shown as box-and-whisker with median (middle line), 25th–75th percentiles (box) and min-max values (whiskers), Tukey-corrected Student’s *t*-test.

KO mice injected with AAV8-Alb-*mbIl11ra1* re-expressed IL11RA1 on hepatocytes and KO mice injected with AAV8-Alb-*sIl11ra1* had increased expression of sIL11RA1 in both the liver and the periphery (**Fig. 6B; Supplementary Fig. 7B, 8B, 8C**). As expected, wild-type mice receiving control AAV8 constructs (AAV8-Alb-Null) on NC had normal livers and, when on either HFMCD diet or WDF, developed steatosis, inflammation and liver damage (**Fig. 6C-J; Supplementary Fig. 7C, 7D, 8D-K**). KO mice injected with control virus and fed either HFMCD or WDF diets were protected from NASH phenotypes, although protection from NASH with germline global deletion of *Il11ra1* was not as strong as seen in the CKOs.

Restoration of IL11 *cis-*signaling in KO mice using mbIl11ra1 recapitulated hepatic steatosis and inflammation that was evident from gross morphology to molecular patterns of gene expression and signaling (**Fig. 6C-6K, Supplementary Fig. 7C, 7D, 8D-L**). However, hepatic collagen content and fibrotic gene expression was not restored (**Fig. 6C, I-J; Supplementary Fig. 7D, 8D, 8I, 8K**) as IL11 signaling in HSCs, important for HSC-to-myofibroblast transformation [7], is unaffected by the albumin-driven Il11ra1 expression (*i.e.* HSCs remain deleted for *Il11ra1* in this model).

In contrast, expression of the sIL11RA in hepatocytes of KOs, which would theoretically activate *trans-*signaling, had no effect despite very high IL11 levels (**Figure 3B**) and mice remained protected from NASH (**Fig. 6C-J; Supplementary Fig. 7C, 7D, 8D-K**). Signaling changes were consistent in that mIL11RA expression restored ERK and JNK activation in KOs on either diet, whereas sIL11RA1 did not (**Fig. 6K; Supplementary Fig. 8L)**. In the WDF model, restoration of hepatocyte-specific IL11 *cis-*signaling in KO mice caused hyperglycemia, hypertriglyceridemia, and hypercholesterolemia but expression of sIl11ra1 did not (**Fig. 6L-N**).

## Discussion

Metabolic liver disease commonly occurs in the context of obesity and type 2 diabetes and manifests initially as NAFLD that can progress to NASH [21,32]. A key underlying pathology in the progression to NASH is “substrate overload”, whereby a nutritional abundance overruns the hepatocyte’s ability to process fat, causing lipotoxicity. Cytokines are key NASH factors secreted from lipotoxic hepatocytes [21] and here we establish IL11 as an important component of the lipotoxic milieu and upstream driver of NAFLD-to-NASH transitions.

A large body of evidence supports the idea that IL6 signaling in the liver is beneficial [16,17,24]. However, a pathogenic role for IL6 *trans-*signaling in hepatic steatosis has also been proposed [29,33]. We found using synthetic constructs that hyperIL11, initiating IL11 *trans-*signaling, is cytotoxic, whereas hyperIL6 is protective in hepatocytes. However, we did not find any evidence for *trans-*signaling in a biologically relevant context *in vitro* or *in vivo*, using both gain- and loss-of-function. We suggest IL6 family member *trans*-signaling plays no role in NASH, which is in agreement with studies outside the liver [19,20].

Here we show the critical importance of IL11 *cis-*signaling in hepatocytes for NASH. This effect was established using both hepatocyte-specific loss-of-function on a wildtype genetic background and also hepatocyte-specific gain-of-function on an *Il11ra1* null background. This overturns the suggestion in the literature that IL11 is protective for hepatocytes based on the use of rhIL11, ineffective in the mouse, in murine models of liver disease [8,10–12]. While restoration of IL11 *cis*-signaling in hepatocytes causes steatohepatitis in KO mice, fibrosis was still prevented, whereas hepatocyte-specific *Il11ra1* deletion protected the CKO mice from both steatohepatitis and fibrosis. This shows that IL11 *cis*-signaling in HSCs is required for liver fibrosis while placing hepatocyte dysfunction upstream of HSC activation in NASH.

Our studies raise questions and have limitations. The published literature suggests IL6R is expressed in hepatocytes [16] and it was surprising that primary human hepatocytes express very little IL6R. This may reflect a strong reliance on transformed hepatocyte-like cells (e.g. HepG2) in earlier studies. The mechanisms underlying the differential activation of ERK/JNK (by IL11) versus STAT3 (IL6) in hepatocytes is perhaps unexpected but consistent across *in vitro* and *in vivo* studies. We documented a surprising effect of inhibiting hepatocyte-specific IL11 *cis*-signaling on weight homeostasis. These various issues require further study.

In conclusion, we propose a new mechanistic model for NAFLD-to-NASH transitions whereby lipid laden hepatocytes secrete IL11 leading to autocrine cell death, paracrine activation of HSCs and secondary inflammatory cell activation and infiltration (**Fig. 7**). Inhibiting IL11 signaling targets an initiating nexus for diet-induced steatohepatitis that impacts subsequent liver fibrosis and inflammation. Hence, therapeutic targeting of IL11-induced lipotoxicity may be beneficial in metabolic liver disease.

**Figure 7.**
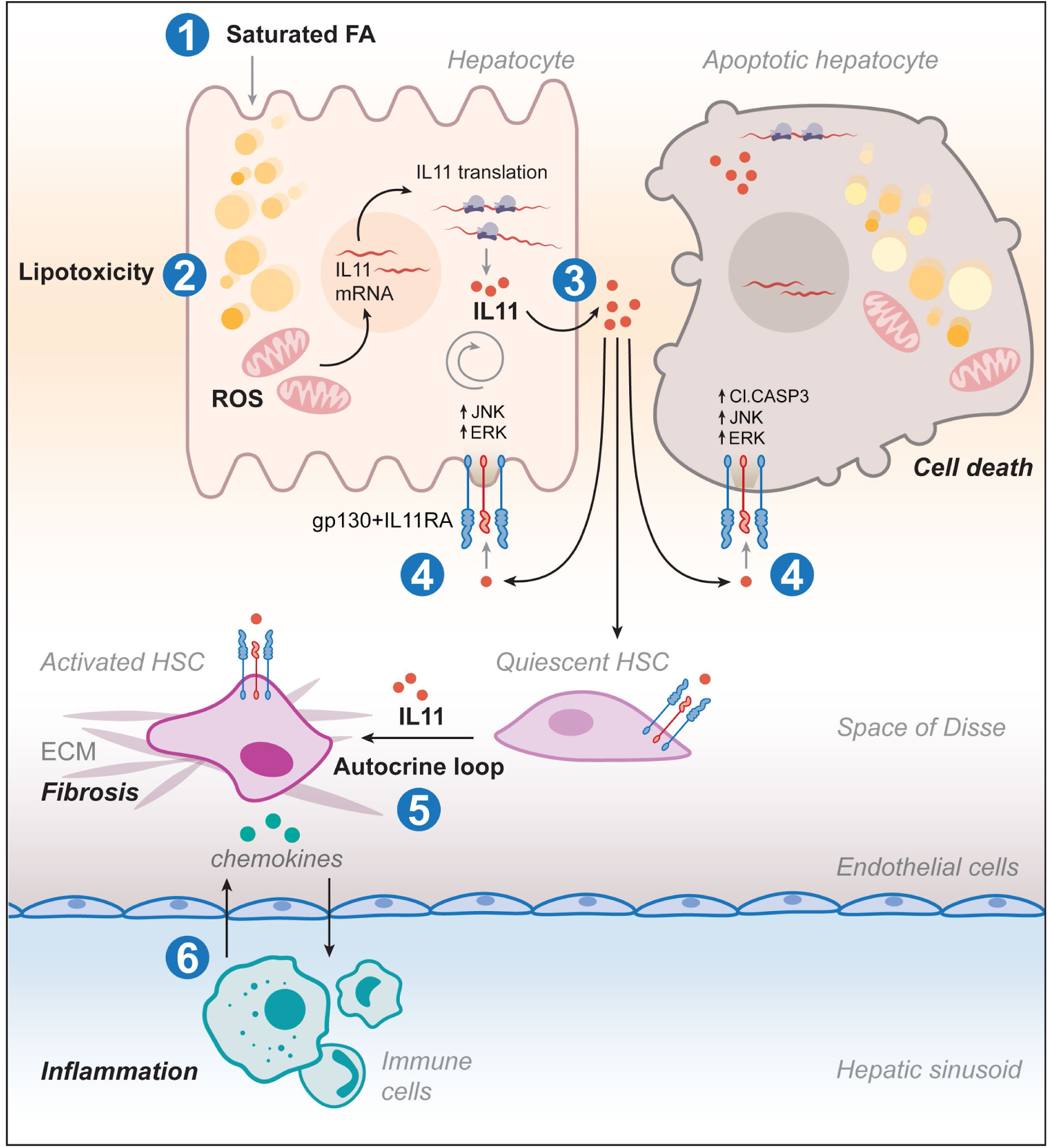
Proposed mechanism for IL11 in NAFLD-to-NASH transition. Excessive lipid accumulation in hepatocytes results in lipotoxicity leading to reactive oxygen species production that triggers IL11 protein secretion. IL11 binds to IL11RA and gp130 on hepatocytes to initiate autocrine ERK, JNK, and Caspase3 activation leading to lipoapoptosis. IL11 also acts in a paracrine fashion to drive transformation of quiescent hepatic stellate cells (HSCs) to become activated myofibroblasts. Cytokines and chemokines released from lipotoxic hepatocytes and HSCs activate and recruit immune cells causing inflammation and progression from NAFLD to NASH.

## Acknowledgements

The authors would like to acknowledge the technical support of B.L.George and S. Lim.

## Data and materials availability

All data are provided in the manuscript or in the supplementary materials.

## References

[1] Schafer S, Viswanathan S, Widjaja AA, Lim W-W, Moreno-Moral A, DeLaughter DM, et al. IL-11 is a crucial determinant of cardiovascular fibrosis. Nature 2017;552:110–5.

[2] Ng B, Dong J, D’Agostino G, Viswanathan S, Widjaja AA, Lim W-W, et al. Interleukin-11 is a therapeutic target in idiopathic pulmonary fibrosis. Sci Transl Med 2019;11. https://doi.org/10.1126/scitranslmed.aaw1237.

[3] Lim W-W, Ng B, Widjaja A, Xie C, Su L, Ko N, et al. Transgenic interleukin 11 expression causes cross-tissue fibro-inflammation and an inflammatory bowel phenotype in mice. PLOS ONE 2020;15:e0227505.https://doi.org/10.1371/journal.pone.0227505.

[4] Cook SA, Schafer S. Hiding in Plain Sight: Interleukin-11 Emerges as a Master Regulator of Fibrosis, Tissue Integrity, and Stromal Inflammation. Annu Rev Med 2020;71:263–76.

[5] Bigaeva E, Gore E, Simon E, Zwick M, Oldenburger A, de Jong KP, et al. Transcriptomic characterization of culture-associated changes in murine and human precision-cut tissue slices. Arch Toxicol 2019. https://doi.org/10.1007/s00204-019-02611-6.

[6] Widjaja AA, Dong J, Adami E, Viswanathan S, Ng B, Singh BK, et al. Redefining Interleukin 11 as a regeneration-limiting hepatotoxin n.d. https://doi.org/10.1101/830018.

[7] Widjaja AA, Singh BK, Adami E, Viswanathan S, Dong J, D’Agostino GA, et al. Inhibiting Interleukin 11 Signaling Reduces Hepatocyte Death and Liver Fibrosis, Inflammation, and Steatosis in Mouse Models of Non-Alcoholic Steatohepatitis. Gastroenterology 2019. https://doi.org/10.1053/j.gastro.2019.05.002.

[8] Nishina T, Komazawa-Sakon S, Yanaka S, Piao X, Zheng D-M, Piao J-H, et al. Interleukin-11 links oxidative stress and compensatory proliferation. Sci Signal 2012;5:ra5.

[9] Bozza M, Bliss JL, Maylor R, Erickson J, Donnelly L, Bouchard P, et al. Interleukin-11 reduces T-cell-dependent experimental liver injury in mice. Hepatology 1999;30:1441–7.

[10] Trepicchio WL, Bozza M, Bouchard P, Dorner AJ. Protective effect of rhIL-11 in a murine model of acetaminophen-induced hepatotoxicity. Toxicol Pathol 2001;29:242–9.

[11] Maeshima K, Takahashi T, Nakahira K, Shimizu H, Fujii H, Katayama H, et al. A protective role of interleukin 11 on hepatic injury in acute endotoxemia. Shock 2004;21:134–8.

[12] Zhu M, Lu B, Cao Q, Wu Z, Xu Z, Li W, et al. IL-11 Attenuates Liver Ischemia/Reperfusion Injury (IRI) through STAT3 Signaling Pathway in Mice. PLoS One 2015;10:e0126296.

[13] Yu J, Feng Z, Tan L, Pu L, Kong L. Interleukin-11 protects mouse liver from warm ischemia/reperfusion (WI/Rp) injury. Clin Res Hepatol Gastroenterol 2016;40:562–70.

[14] Wuestefeld T, Klein C, Streetz KL, Betz U, Lauber J, Buer J, et al. Interleukin-6/glycoprotein 130-dependent pathways are protective during liver regeneration. J Biol Chem 2003;278:11281–8.

[15] Klein C, Wüstefeld T, Assmus U, Roskams T, Rose-John S, Müller M, et al. The IL-6-gp130-STAT3 pathway in hepatocytes triggers liver protection in T cell-mediated liver injury. J Clin Invest 2005;115:860–9.

[16] Schmidt-Arras D, Rose-John S. IL-6 pathway in the liver: From physiopathology to therapy. J Hepatol 2016;64:1403–15.

[17] Kroy DC, Beraza N, Tschaharganeh DF, Sander LE, Erschfeld S, Giebeler A, et al. Lack of interleukin-6/glycoprotein 130/signal transducers and activators of transcription-3 signaling in hepatocytes predisposes to liver steatosis and injury in mice. Hepatology 2010;51:463–73.

[18] Matthews VB, Allen TL, Risis S, Chan MHS, Henstridge DC, Watson N, et al. Interleukin-6-deficient mice develop hepatic inflammation and systemic insulin resistance. Diabetologia 2010;53:2431–41.

[19] Balic JJ, Garbers C, Rose-John S, Yu L, Jenkins BJ. Interleukin-11-driven gastric tumourigenesis is independent of trans-signalling. Cytokine 2017;92:118–23.

[20] Agthe M, Garbers Y, Putoczki T, Garbers C. Interleukin-11 classic but not trans-signaling is essential for fertility in mice. Placenta 2017;57:13–6.

[21] Friedman SL, Neuschwander-Tetri BA, Rinella M, Sanyal AJ. Mechanisms of NAFLD development and therapeutic strategies. Nat Med 2018. https://doi.org/10.1038/s41591-018-0104-9.

[22] Farrell GC, Haczeyni F, Chitturi S. Pathogenesis of NASH: How Metabolic Complications of Overnutrition Favour Lipotoxicity and Pro-Inflammatory Fatty Liver Disease. Adv Exp Med Biol 2018;1061:19–44.

[23] Kakisaka K, Cazanave SC, Fingas CD, Guicciardi ME, Bronk SF, Werneburg NW, et al. Mechanisms of lysophosphatidylcholine-induced hepatocyte lipoapoptosis. Am J Physiol Gastrointest Liver Physiol 2012;302:G77–84.

[24] Yamaguchi K, Itoh Y, Yokomizo C, Nishimura T, Niimi T, Fujii H, et al. Blockade of interleukin-6 signaling enhances hepatic steatosis but improves liver injury in methionine choline-deficient diet-fed mice. Lab Invest 2010;90:1169–78.

[25] Alsabeeh N, Chausse B, Kakimoto PA, Kowaltowski AJ, Shirihai O. Cell culture models of fatty acid overload: Problems and solutions. Biochim Biophys Acta Mol Cell Biol Lipids 2018;1863:143–51.

[26] Ng B, Dong J, Viswanathan S, Widjaja AA, Paleja BS, Adami E, et al. Fibroblast-specific IL11 signaling is required for lung fibrosis and inflammation n.d. https://doi.org/10.1101/801852.

[27] Kleinfeld AM, Prothro D, Brown DL, Davis RC, Richieri GV, DeMaria A. Increases in serum unbound free fatty acid levels following coronary angioplasty. Am J Cardiol 1996;78:1350–4.

[28] Bettaieb A, Jiang JX, Sasaki Y, Chao T-I, Kiss Z, Chen X, et al. Hepatocyte Nicotinamide Adenine Dinucleotide Phosphate Reduced Oxidase 4 Regulates Stress Signaling, Fibrosis, and Insulin Sensitivity During Development of Steatohepatitis in Mice. Gastroenterology 2015;149:468–80.e10.

[29] Kammoun HL, Allen TL, Henstridge DC, Kraakman MJ, Peijs L, Rose-John S, et al. Over-expressing the soluble gp130-Fc does not ameliorate methionine and choline deficient diet-induced non alcoholic steatohepatitis in mice. PLoS One 2017;12:e0179099.

[30] Kraakman MJ, Kammoun HL, Allen TL, Deswaerte V, Henstridge DC, Estevez E, et al. Blocking IL-6 trans-signaling prevents high-fat diet-induced adipose tissue macrophage recruitment but does not improve insulin resistance. Cell Metab 2015;21:403–16.

[31] Stephenson K, Kennedy L, Hargrove L, Demieville J, Thomson J, Alpini G, et al. Updates on Dietary Models of Nonalcoholic Fatty Liver Disease: Current Studies and Insights. Gene Expr 2018;18:5–17.

[32] Sanyal AJ. Past, present and future perspectives in nonalcoholic fatty liver disease. Nat Rev Gastroenterol Hepatol 2019;16:377–86.

[33] Wieckowska A, Papouchado BG, Li Z, Lopez R, Zein NN, Feldstein AE. Increased hepatic and circulating interleukin-6 levels in human nonalcoholic steatohepatitis. Am J Gastroenterol 2008;103:1372–9.

