## Supplementary Data for "Autocrine IL11 cis-signaling in hepatocytes is an initiating nexus between lipotoxicity and non-alcoholic steatohepatitis"

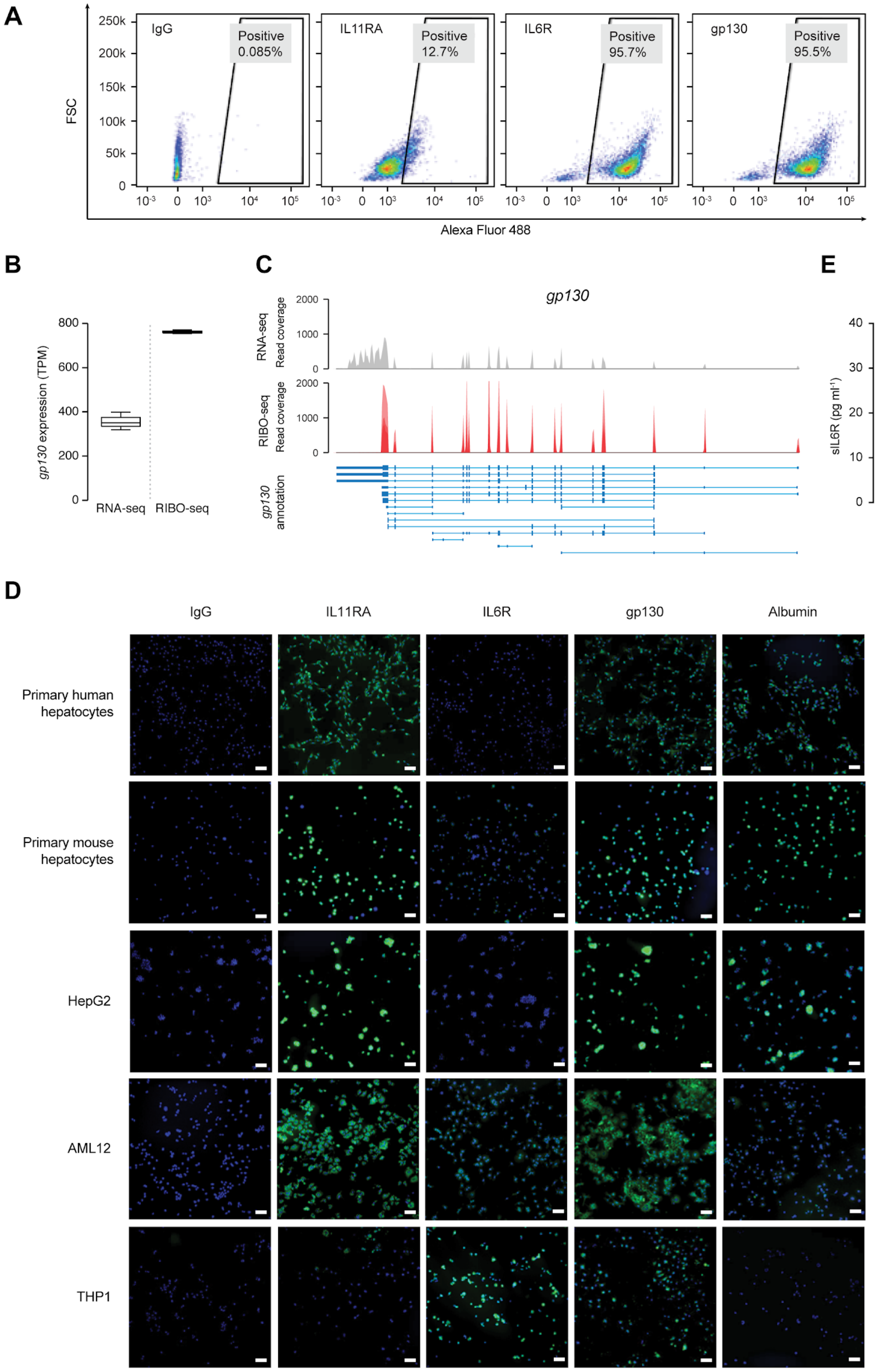
A
IgG
Positive 0.085%
IL11RA
Positive 12.7%
IL6R
Positive 95.7%
gp130
Positive 95.5%
FSC
Alexa Fluor 488
B
gp130 expression (TPM)
RNA-seq
RIBO-seq
C
gp130
RNA-seq Read coverage
RIBO-seq Read coverage
gp130 annotation
D
IgG
IL11RA
IL6R
gp130
Albumin
Primary human hepatocytes
Primary mouse hepatocytes
HepG2
AML12
THP1
E
sIL6R (pg ml-1)
Primary human hepatocytes

**Supplementary Figure 1. Primary human hepatocytes express IL11RA but not IL6R.**

(A) Representative FSC plots of IL11RA, IL6R, and gp130 staining on activated THP-1 cells. (B) *gp130* transcripts in primary human hepatocytes based on RNA-seq and Ribo-seq (TPM). (C) Read coverage of *gp130* transcripts based on RNA-seq (gray) and Ribo-seq (red) of primary human hepatocytes (n=3). (D) Immunofluorescence images (scale bars, 100 μm) of IL11RA, IL6R, gp130, and Albumin expression in primary human hepatocytes and activated THP-1 cells. (E) The levels of soluble IL6R in the hepatocyte media at basal. (B) Data are shown as box-and-whisker with median (middle line), 25th–75th percentiles (box) and min-max values (whiskers); (E) data are shown as mean ± SEM; (B, E) Tukey-corrected Student's *t*-test.

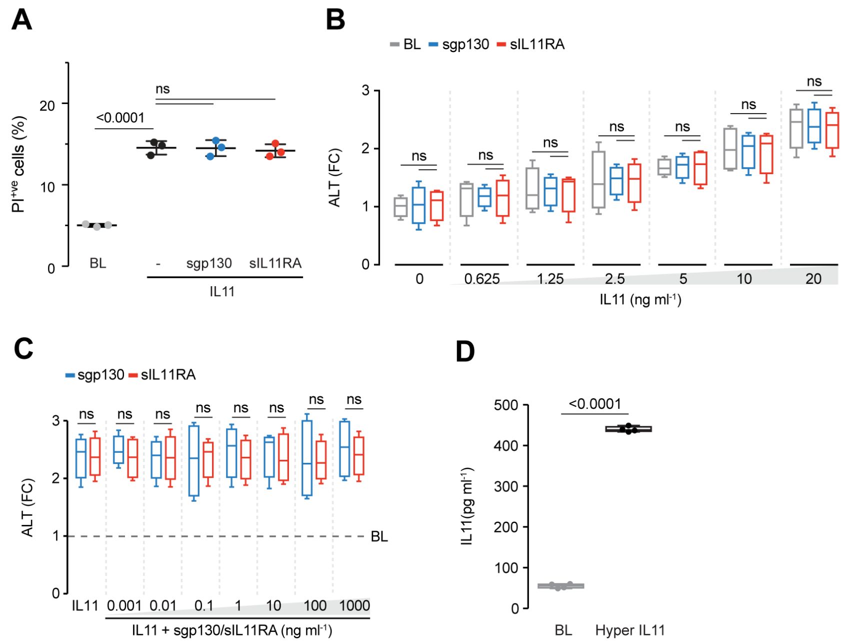


**Supplementary Figure 2. No evidence to support the existence of physiologically occurring IL11 *trans*-signaling in hepatocytes.**

(A) Quantification of PI staining on IL11-stimulated primary human hepatocytes (PI$^{+ve}$ cells) in the presence of sgp130 or sIL11RA. (B) Dose-dependent effect of increasing concentration of IL11 in the presence of 1 µg/ml of sgp130 or sIL11RA on ALT levels secreted by primary human hepatocytes. (C) Dose-dependent effect of increasing concentration of either sgp130 or sIL11RA on IL11-induced ALT secretion. (D) ELISA of IL11 expression after hyperIL11 treatment (20 ng/ml). (A) Data are shown as mean ± SEM; (B-D) data are shown as box-and-whisker with median (middle line), 25th–75th percentiles (box) and min-max values (whiskers); (A-C) Tukey-corrected Student's *t*-test; (D) two-tailed Student's *t*-test

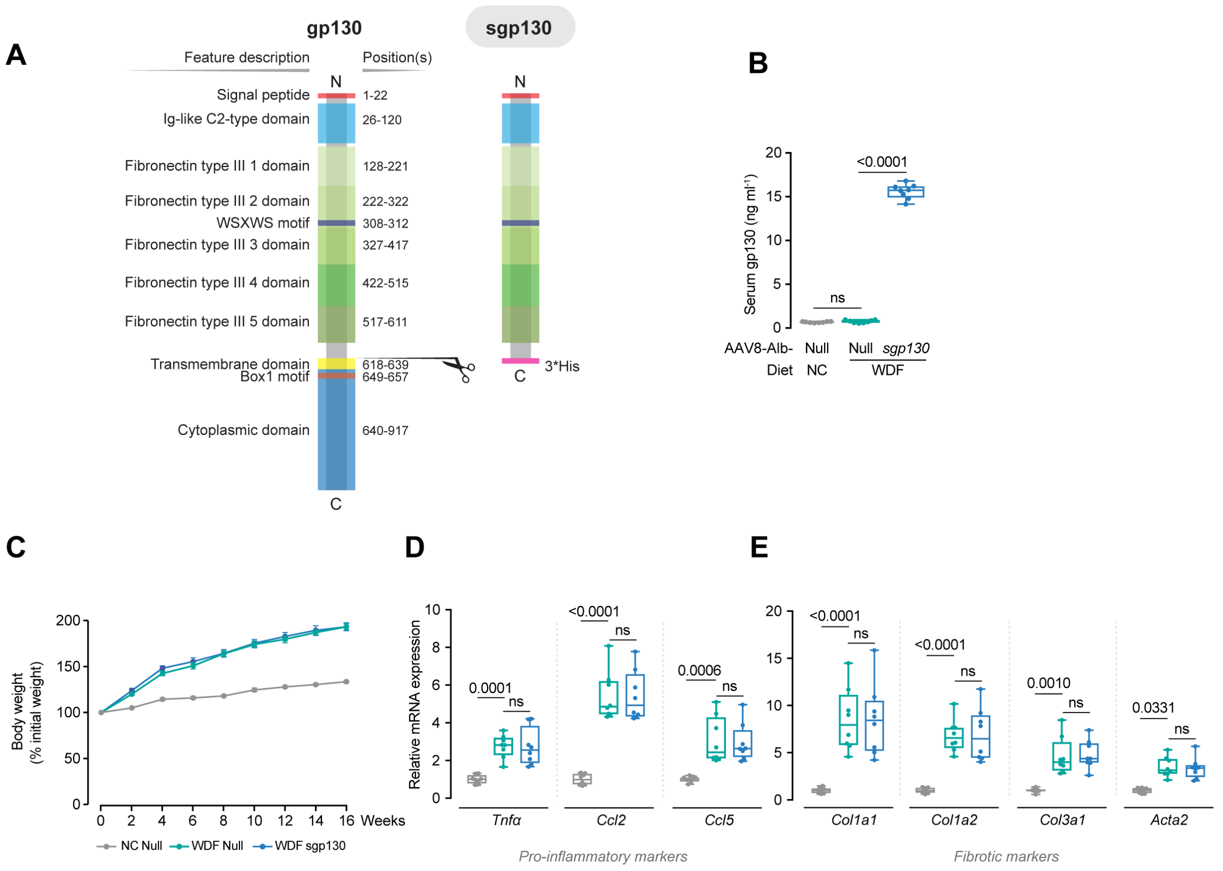


**Supplementary Figure 3. Sgp130 expression does not protect mice from WDF-induced liver and obesity phenotypes.**

(A) Schematic of gp130 protein domain structure and its amino acid position (left) and the domains that were used to construct sgp130 in this study (right). (B-E) Data for WDF-sgp130 *in vivo* experiments as shown in **Figure 3A**. (B) Serum gp130 levels in NC-fed control mice and WDF-fed AAV8-Alb-Null- and AAV8-Alb-sgp130-injected mice. (C) Effect of 16 weeks of WDF on body weight of AAV8-Alb-Null- and AAV8-Alb-sgp130-injected mice. Data are shown as mean ± SEM. (D and E) Hepatic mRNA expression of (D) pro-inflammatory markers (*Tnf*□, *Ccl2*, *Ccl5*) and (E) fibrosis markers (*Col1a1, Col1a2, Col3a1, Acta2*) as shown in **Figure 3O**. (B, D-E) Data are shown as box-and-whisker with median (middle line), 25th–75th percentiles (box) and min-max values (whiskers), Tukey-corrected Student's *t*-test.

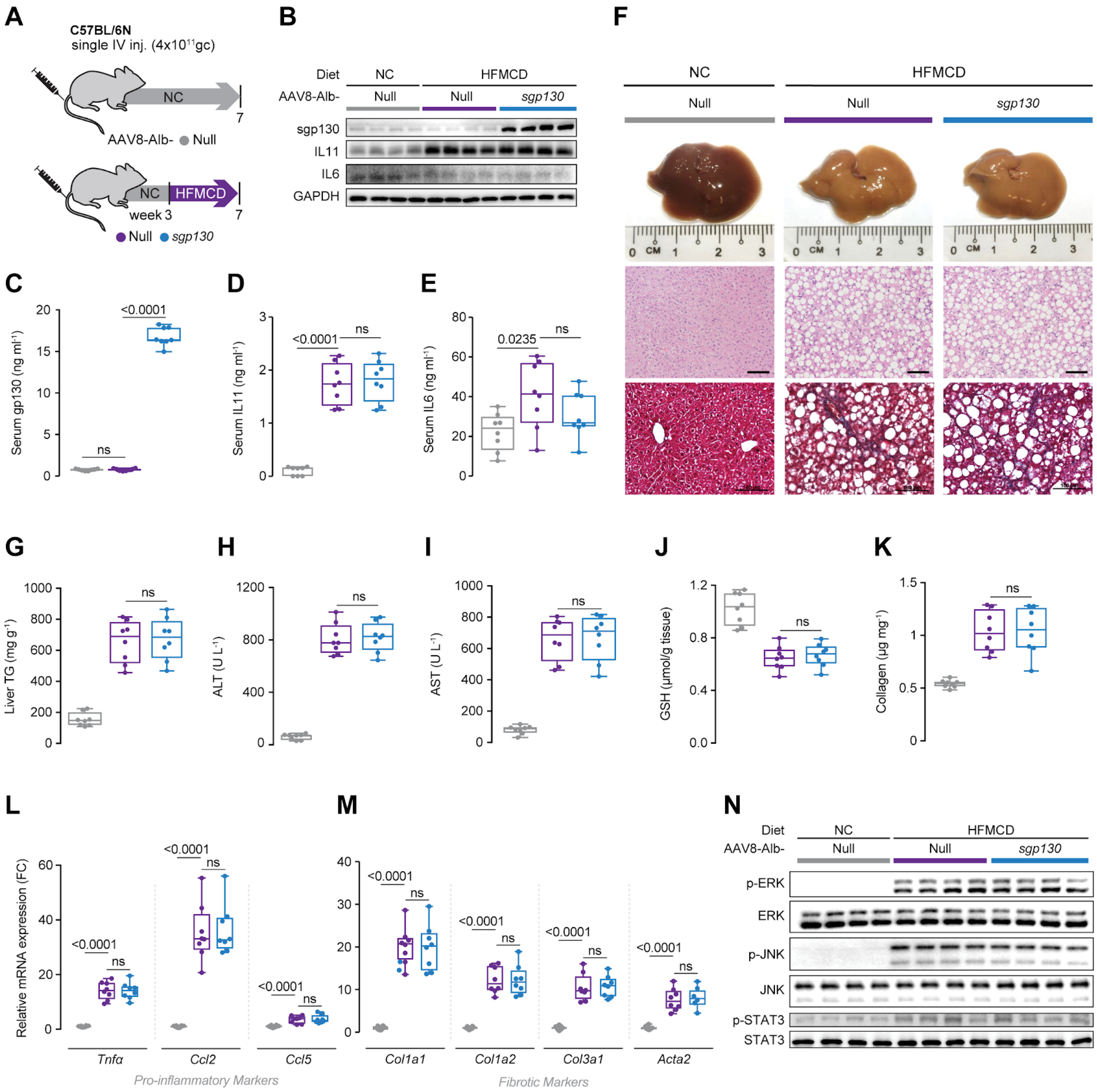


**Supplementary Figure 4. Inhibition of putative *trans*-signaling of IL6 family members has no effect on NASH phenotypes in mice on HFMCD diet.**
(A) Schematic of mice with hepatocyte-specific expression of sgp130 in mice on HFMCD diets for data shown in (B-N). Mice were intravenously injected with either AAV8-Alb-Null or AAV8-Alb-sgp130 and fed HFMCD for 4 weeks. (B) Western blots showing hepatic levels of sgp130, IL11 and IL6 with GAPDH shown as internal control. (C) Serum gp130 levels. (D) Serum IL11 levels. (E) Serum IL6 levels. (F) Representative gross anatomy, H&E-stained (scale bars, 50 μm) and Masson's Trichrome (scale bars, 100 μm) images of livers. (G) Hepatic triglycerides content. (H) Serum ALT levels. (I) Serum AST levels. (J) Hepatic GSH content. (K) Hepatic collagen levels. (L and M) Hepatic mRNA expression of (L) pro-inflammatory markers (*Tnf*[illegible],

*Ccl2*, *Ccl5*) and (M) fibrosis markers (*Col1a1, Col1a2, Col3a1, Acta2*). (N) Western blots of hepatic p-ERK, ERK, p-JNK, JNK, p-STAT3, STAT3. (C-E, G-M) Data are shown as box-and-whisker with median (middle line), 25th–75th percentiles (box) and min-max values (whiskers), Tukey-corrected Student's *t*-test.

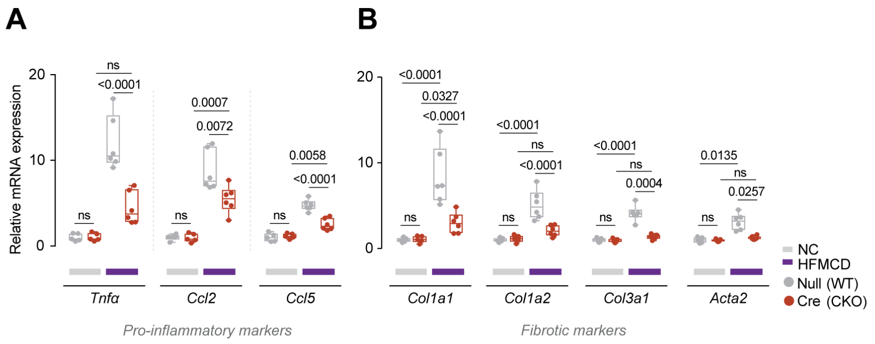


**Supplementary Figure 5. Mice with hepatocyte-specific deletion of *Il11ra1* are protected from HFMCD-induced gene dysregulation.**

(A and B) Hepatic mRNA expression of (A) pro-inflammatory markers (*Tnf*□, *Ccl2*, *Ccl5*) and (B) fibrotic markers (*Col1a1, Col1a2, Col3a1, Acta2*) from control and CKO mice on NC and HFMCD diet as shown in **Figure 4J**. (A-B) Data are shown as box-and-whisker with median (middle line), 25th–75th percentiles (box) and min-max values (whiskers), Sidak-corrected Student's *t*-test.

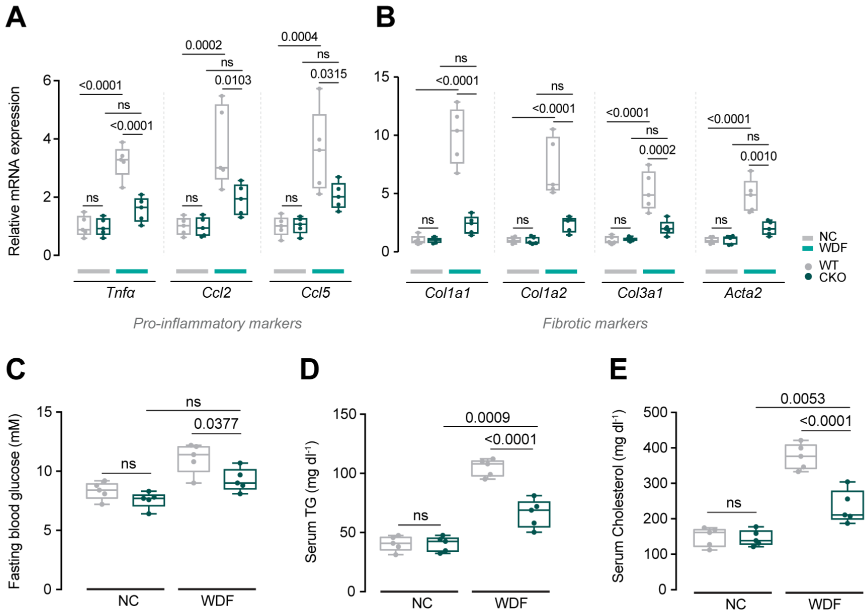


**Supplementary Figure 6. Hepatocyte-specific *Il11ra1* deleted mice are protected from WDF-induced NASH phenotypes.**

(A-E) Data for control and CKO mice on NC and WDF diet as shown in **Figure 5A**. (A and B) Hepatic mRNA expression of (A) pro-inflammatory markers (*Tnf*[illegible], *Ccl2*, *Ccl5*) and (B) fibrotic markers (*Col1a1, Col1a2, Col3a1, Acta2*) as shown in **Figure 5L**. (C) Fasting blood glucose levels. (D) Serum triglycerides levels. (E) Serum cholesterol levels. (A-E) Data are shown as box-and-whisker with median (middle line), 25th–75th percentiles (box) and min-max values (whiskers), Sidak-corrected Student’s *t*-test.

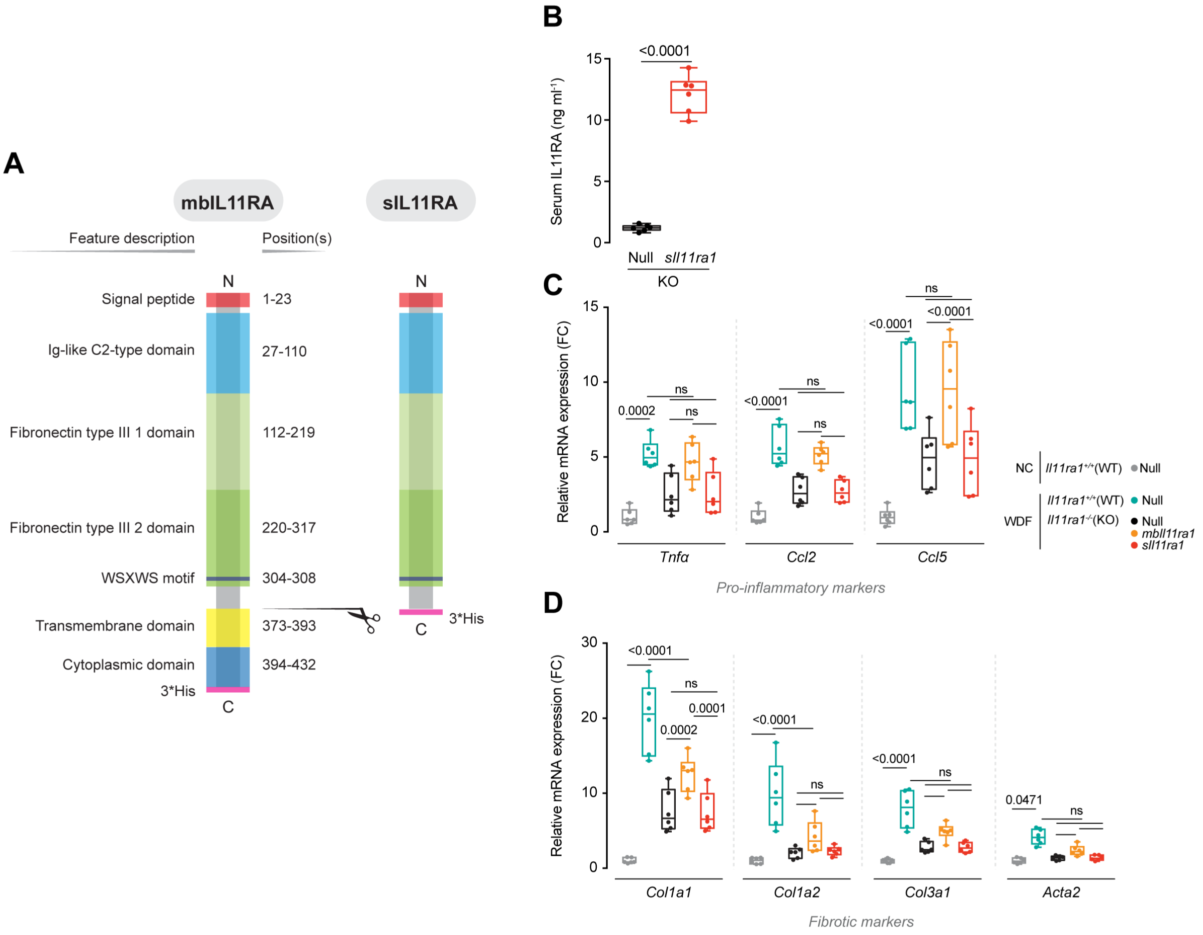


**Supplementary Figure 7. Hepatocyte-specific IL11 *cis*-signaling but not IL11 *trans*-signaling drives WDF-induced steatohepatitis in mice.**
(A) Schematic of full-length membrane-bound IL11RA protein domain structure and its amino acid position (left) and the domains that were used to construct soluble IL11RA (right). (B-D) Data for WDF feeding regimen on *Il11ra1*$^{+/+}$ (WT) mice and mice globally deleted for *Il11ra* (*Il11ra1*$^{-/-}$ ;KO mice) that had been injected with AAV8-Alb-Null, AAV8-Alb-mbIl11ra1 (full length membrane-bound Il11ra1) or AAV8-Alb-sIl11ra1 (soluble form of Il11ra1) as illustrated in **Figure 6A.** (B) Serum IL11RA levels in AAV8-Alb-Null and AAV8-Alb-sIl11ra1-injected KO mice on WDF. (C and D) Hepatic mRNA expression of (C) pro-inflammatory markers (*Tnf*□, *Ccl2*, *Ccl5*) and (D) fibrotic markers (*Col1a1, Col1a2, Col3a1, Acta2*). (B-D) Data are shown as box-and-whisker with median (middle line), 25th–75th percentiles (box) and min-max values (whiskers), Tukey-corrected Student's *t*-test.

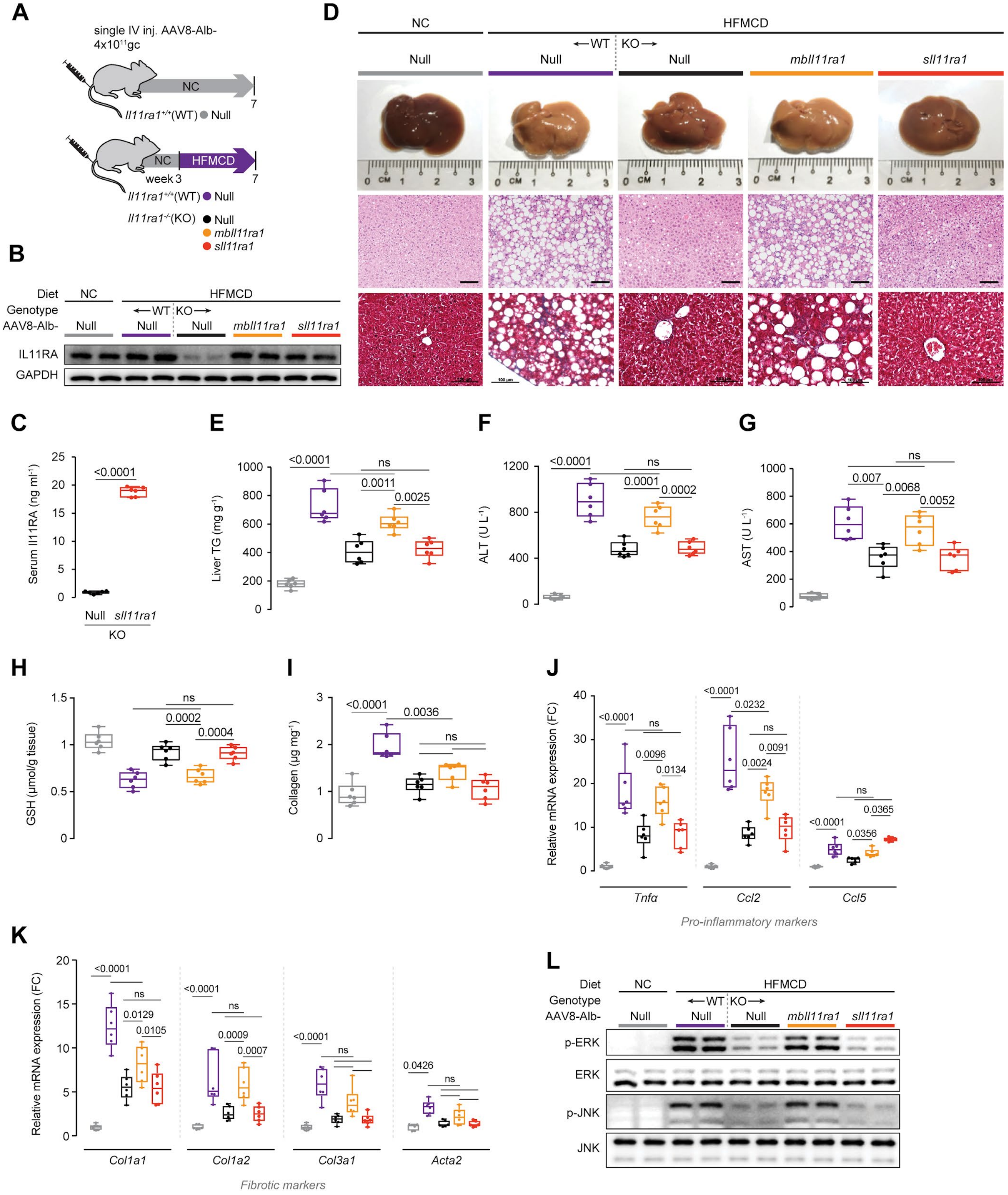


**Supplementary Figure 8. Hepatocyte-specific IL11 *cis*-signaling but not IL11 *trans*-signaling drives steatohepatitis in mice on a HFMCD.**

(A) Schematic of HFMCD-fed WT and KO mice for experiments shown in (B-L). KO mice were intravenously injected with either AAV8-Alb-Null, AAV8-Alb-mbIl11ra1 or AAV8-ALB-sIl11ra1; WT mice received AAV8-Alb-Null as control. Three weeks following virus administration, mice were started on HFMCD feeding for 4 weeks. (B) Western blots showing hepatic levels of IL11RA and GAPDH. (C) Serum IL11RA levels. (D) Representative gross anatomy, H&E-stained (scale bars, 50 μm) and Masson's Trichrome (scale bars, 100 μm) images of livers. (E) Hepatic triglycerides content. (F) Serum ALT levels. (G) Serum AST levels. (H) Hepatic GSH levels. (I) Hepatic collagen content. (J and K) Hepatic mRNA expression of (J) pro-inflammatory markers (*Tnf*□, *Ccl2*, *Ccl5*) and (K) fibrotic markers (*Col1a1, Col1a2, Col3a1, Acta2*). (M) Western blots showing activation status of hepatic ERK and JNK. (C, E-K) Data are shown as box-and-whisker with median (middle line), 25th–75th percentiles (box) and min-max values (whiskers), Tukey-corrected Student's *t*-test.

### Supplementary Materials and Methods

#### *AAV8 vectors*

All Adeno-associated virus serotype 8 (AAV8) vectors used in this study were synthesized by Vector Biolabs. AAV8 vector carrying a mouse membrane-bound *Il11ra1* cDNA (NCBI accession number: BC069984), a mouse soluble *Il11ra1* cDNA , and a mouse soluble *gp130* cDNA driven by Albumin (*Alb*) promoter is referred to as AAV8-*Alb-mbIl11ra1*, AAV8-*Alb-sIl11ra1*, and AAV8-*Alb-sgp130*, respectively. AAV8-*Alb-sgp130* and AAV8-*Alb-sIl11ra1* were constructed by removing the transmembrane and cytoplasmic regions of mouse *gp130* sequence (NCBI accession number: BC058679) and mouse *Il11ra1* sequence, respectively. AAV8-Null vector was used as vector control. To specifically delete *Il11ra1* in *Albumin*-expressing cells, AAV8-*Alb-iCre* vector was injected to mice homozygous for LoxP-flanked *Il11ra1* alleles (*Il11ra1*$^{loxP/loxP}$ mice).

#### *Antibodies*

Albumin (ab207327, Abcam), Alexa Fluor 488 secondary antibody (ab150077, Abcam), Cleaved Caspase-3 (9664, CST), Caspase-3 (9662, CST), p-ERK1/2 (4370, CST), ERK1/2 (4695, CST), FASN (ab128856, Abcam), GAPDH (2118, CST), gp130 (PA5-28932, Thermo Fisher), gp130 (extracellular, PA5-77476, Thermo Fisher), IL6 (AF506, R&D systems), IL6R (flow cytometry, ab222101, Abcam), IL6R (human, IHC and IF, MA1-80456, Thermo Fisher), IL6R (mouse, IF, ab83053, Abcam), IL11 (Aldevron), IL11RA (IHC, IF, flow cytometry, ab125015, Abcam), IL11RA (western blot, 130920, Santa Cruz), p-JNK (4668, CST), JNK (9258, CST), NOX4 (110-58849, Novus Biologicals), p-STAT3 (4113, CST), STAT3 (4904, CST), mouse HRP (7076, CST), rabbit HRP (7074, CST), rat HRP (31470, Santa Cruz).

#### *Recombinant proteins*

Commercial recombinant proteins: Human hyperIL6 (IL6R:IL6 fusion protein, 8954-SR, R&D systems), human soluble gp130 Fc (671-GP-100, R&D systems), human IL11RA (8895-MR-050, R&D systems).
Custom recombinant proteins: Human IL11 (UniProtKB:P20809, Genscript). Human hyperIL11 (IL11RA:IL11 fusion protein), which mimics the *trans*-signalling complex, was constructed using a fragment of IL11RA (amino acid residues 1–317; UniProtKB: Q14626) and IL11 (amino acid residues 22–199, UniProtKB: P20809) with a 20 amino acid linker (GPAGQSGGGGGSGGGSGGGSV) (Schafer et al., 2017).

#### *Chemicals*

Palmitate (P5585, Sigma), Paraformaldehyde (PFA, 28908; Thermo Fisher), phorbol 12-myristate 13-acetate (PMA, P1585, Sigma), Triton X-100 (T8787, Sigma), and 4’,6-diamidino-2-phenylindole (D1306; Thermo Fisher).

***Oil Red O Staining***

Primary human hepatocytes were seeded on 8-well chamber slides (1X$10^4$ cells/well) Following 24 hours of palmitate treatment, cells were fixed in 10% PFA for 30 minutes, washed with distilled water, and incubated with 60% (v/v) isopropyl alcohol for 5 minutes. Cells were then stained with Oil Red O Solution for 30 minutes and washed with distilled water prior to imaging with bright field microscope (BX53, Olympus). The lipid droplets were identified by their red staining.

***Reactive Oxygen Species (ROS) Detection***

Primary human hepatocytes were seeded on 8-well chamber slides (1X$10^4$ cells/well). For this experiment, cells were not serum-starved prior to palmitate treatment. 24 hours following palmitate stimulation, cells were washed, incubated with 25 µM of DCFDA solution (ab113851, Abcam) for 45 minutes at 37°C in the dark, and rinsed with dilution buffer according to the manufacturer's protocol. Live cells with positive DCF staining were imaged with filter set appropriate for fluorescein (FITC) using a fluorescence microscope (Leica).

***RNA-sequencing (RNA-seq) and Ribosome profiling (Ribo-seq)***

RNA-seq and Ribo-Seq library preparations were performed as previously described (Chothani et al., 2019).

*Generation of RNA-seq libraries*

Total RNA was extracted from human hepatocytes using RNeasy columns (Qiagen). RNA was quantified using a Qubit RNA High-Sensitivity Assay kit (Life Technologies) and its quality was assessed on the basis of their RNA integrity number using the LabChip GX RNA Assay Reagent Kit (Perkin Elmer). TruSeq Stranded mRNA Library Preparation kit (Illumina) was used to measure transcript abundance following standard instructions from the manufacturer.

*Generation of Ribo-seq libraries*

Hepatocytes were grown to 90% confluence in a 10cm culture dish and lysed in 1 mL cold lysis buffer (formulation as in TruSeq® Ribo Profile Mammalian Kit, RPHMR12126, Illumina) supplemented with 0.1 mg/mL cycloheximide. Homogenized and cleared lysates were then footprinted with Truseq Nuclease (Illumina) according to the manufacturer’s instructions. Ribosomes were purified using Illustra Sephacryl S400 columns (GE Healthcare), and the protected RNA fragments were extracted with a standard phenol:chloroform:isoamylalcohol technique. Following ribosomal RNA removal (Mammalian RiboZero Magnetic Gold, Illumina), sequencing libraries were then prepared out of the footprinted RNA by using TruSeq® Ribo Profile Mammalian Kit according to the manufacturer’s protocol.

The final RNA-seq and ribosome profiling libraries were quantified using KAPA library quantification kits (KAPA Biosystems) on a StepOnePlus Real-Time PCR system (Applied Biosystems) according to the manufacturer's protocol. The quality and average fragment size of the final libraries were determined using a LabChip GX DNA High Sensitivity Reagent Kit (Perkin Elmer). Libraries with unique indexes were pooled and sequenced on a NextSeq 500 benchtop sequencer (Illumina) using NextSeq 500 High Output v2 kit and paired-end 75-bp sequencing chemistry.

*Data processing and analyses for RNA-sequencing and Ribosome profiling*

Raw sequencing data were demultiplexed with bcl2fastq *V2.19.0.316* and the adaptors were trimmed using *Trimmomatic (Bolger et al., 2014) V0.36*, retaining reads longer than 20 nt post-clipping. Ribo-seq reads were aligned using bowtie (Langmead et al., 2009) to known mtRNA, rRNA and tRNA sequences (RNACentral(The RNAcentral Consortium, 2017), release 5.0) and only unaligned reads were retained as Ribosome protected fragments (RPFs). Alignment to the human genome (hg38) was carried out using STAR (Dobin et al., 2012). Gene expression was quantified on the CDS (coding sequence) regions for Ribo-seq and exonic regions for RNA-seq using uniquely mapped reads (Ensembl database release GRCh38 v86) with feature counts (Liao et al., 2014). TPM was calculated and visualized using boxplot to compare baseline expression of IL11RA (ENSG00000137070), IL6R (ENSG00000160712), and gp130 (ENSG00000134352). Read coverage using Ribo-seq and RNA-seq reads for IL11RA, IL6R and gp130 was visualized using Gviz R package (Hahne and Ivanek, 2016) with strand specific alignment files.

***Colorimetric assays***

Alanine Aminotransferase (ALT) activity in the cell culture supernatant and mouse serum was measured using ALT Activity Assay Kit (ab105134, Abcam). Liver Glutathione (GSH) levels were measured using Glutathione Colorimetric Detection Kit (EIAGSHC, Thermo Fisher). Total hydroxyproline content in mouse livers was measured using Quickzyme Total Collagen assay kit (QZBtotco15, Quickzyme Biosciences). The levels of serum and liver triglycerides were measured using Triglyceride Assay Kit (ab65336, Abcam). Mouse serum levels of Aspartate Aminotransferase (AST) and cholesterol were measured using AST Assay Kit (ab105135, Abcam) and Cholesterol Assay Kit (ab65390; Abcam), respectively. All colorimetric assays were performed according to the manufacturer's protocol.

***Enzyme-linked immunosorbent assay (ELISA)***

The levels of gp130 in mouse serum were quantified using Mouse gp130 DuoSet ELISA (DY468, R&D systems) according to the manufacturer's protocol.

***RT-qPCR***

Total RNA was extracted from snap-frozen liver tissues using Trizol (Invitrogen) and RNeasy Mini Kit (Qiagen). PCR amplifications were performed using iScript cDNA Synthesis Kit (Biorad). Gene expression was analyzed in duplicate by TaqMan (Applied Biosystems) or SYBR green (Qiagen) technology using StepOnePlus (Applied Biosystem) over 40 cycles. Expression data were normalized to *GAPDH* mRNA expression and fold change was calculated using $2^{-\Delta\Delta Ct}$ method. The sequences of specific TaqMan probes and SYBR green primers are available upon request.

***Immunoblotting***
Western blots were carried out on total protein extracts from hepatocytes and liver tissues. Hepatocyte and liver tissue lysates were homogenized in RIPA Lysis and Extraction Buffer (89901, Thermo Scientific) containing protease and phosphatase inhibitors (Roche). Protein lysates were separated by SDS-PAGE and transferred to PVDF membranes. Protein bands were visualized using the ECL detection system (Pierce) with the appropriate secondary antibodies: anti-rabbit HRP or anti-mouse HRP.

***Liver tissue processing and histological analysis***
*Immunohistochemistry (IHC)*
Serial sections of healthy human liver sections (ab4348, Abcam) were immunostained with IL11RA or IL6R antibody. Tissue sections were incubated with primary antibodies overnight and visualized using an ImmPRESS HRP anti-rabbit IgG polymer detection kit (MP-7401, Vector Laboratories) with ImmPACT DAB Peroxidase Substrate (SK-4105, Vector Laboratories).
*H&E and Masson's Trichrome staining*
Mouse liver samples were fixed in 10% neutral formalin, paraffinized, cut into 7µm sections, stained with hematoxylin and eosin (H&E) or Masson's Trichrome according to standard protocol, and examined by light microscopy.

***Statistical analysis***
All statistical analyses were performed using GraphPad Prism software (version 6.07). P values were corrected for multiple testing according to Dunnett's (when several experimental groups were compared to one condition), Tukey (when several conditions were compared to each other within one experiment), Sidak (when several conditions from 2 different genotypes were compared to each other). Analysis for two parameters for comparison of two different groups were performed by two-way ANOVA. The criterion for statistical significance was set at $P < 0.05$.

### References


Alsabeeh, N., Chausse, B., Kakimoto, P.A., Kowaltowski, A.J., and Shirihai, O. (2018). Cell culture models of fatty acid overload: Problems and solutions. Biochim. Biophys. Acta Mol. Cell Biol. Lipids *1863*, 143–151.

Bolger, A.M., Lohse, M., and Usadel, B. (2014). Trimmomatic: a flexible trimmer for Illumina sequence data. Bioinformatics *30*, 2114–2120.

Chothani, S., Schäfer, S., Adami, E., Viswanathan, S., Widjaja, A.A., Langley, S.R., Tan, J., Wang, M., Quaife, N.M., Jian Pua, C., et al. (2019). Widespread Translational Control of Fibrosis in the Human Heart by RNA-Binding Proteins. Circulation *140*, 937–951.

Dobin, A., Davis, C.A., Schlesinger, F., Drenkow, J., Zaleski, C., Jha, S., Batut, P., Chaisson, M., and Gingeras, T.R. (2012). STAR: ultrafast universal RNA-seq aligner. Bioinformatics *29*, 15–21.

Hahne, F., and Ivanek, R. (2016). Visualizing Genomic Data Using Gviz and Bioconductor. In Statistical Genomics, (Humana Press, New York, NY), pp. 335–351.

Kleinfeld, A.M., Prothro, D., Brown, D.L., Davis, R.C., Richieri, G.V., and DeMaria, A. (1996). Increases in serum unbound free fatty acid levels following coronary angioplasty. Am. J. Cardiol. *78*, 1350–1354.

Langmead, B., Trapnell, C., Pop, M., and Salzberg, S.L. (2009). Ultrafast and memory-efficient alignment of short DNA sequences to the human genome. Genome Biol. *10*, R25.

Liao, Y., Smyth, G.K., and Shi, W. (2014). featureCounts: an efficient general purpose program for assigning sequence reads to genomic features. Bioinformatics *30*, 923–930.

Ng, B., Dong, J., Viswanathan, S., Widjaja, A.A., Paleja, B.S., Adami, E., Ko, N.S.J., Wang, M., Lim, S., Tan, J., et al. Fibroblast-specific IL11 signaling is required for lung fibrosis and inflammation.

Schafer, S., Viswanathan, S., Widjaja, A.A., Lim, W.-W., Moreno-Moral, A., DeLaughter, D.M., Ng, B., Patone, G., Chow, K., Khin, E., et al. (2017). IL-11 is a crucial determinant of cardiovascular fibrosis. Nature *552*, 110–115.

The RNAcentral Consortium (2017). RNAcentral: a comprehensive database of non-coding RNA sequences. Nucleic Acids Res. *45*, D128–D134.
